# Maize Kernel Development Stage the Primary Factor in Differential Gene Expression in Response to Two Methods of Field Inoculation with *Aspergillus flavus*

**DOI:** 10.1101/617241

**Authors:** Nancy J. Wahl, Seth C. Murray, Hong-Bin Zhang, Meiping Zhang, C. Michael Dickens, Thomas S. Isakeit

## Abstract

**Background:** Aflatoxins, produced by the fungus *Aspergillus flavus*, often contaminate preharvest maize (*Zea mays* L.) grain under heat and drought stresses, posing serious health hazards to humans and livestock, and resulting in significant costs to identify and dispose of contaminated grain. This study was designed to investigate the changes in differential gene expression during seed morphogenesis and maturation in the “aflatoxin resistant” Argentinian inbred line Tx772 when challenged by the introduction of *A. flavus* through two different methods of ear inoculation; non-wounding (silk channel, used to select Tx772), wounding (side needle) and a non-inoculated control.

**Method and Findings:** Grain maturity had the largest effect on overall RNA-Seq differential gene expression (DGE) as measured by edgeR of the Bioconductor platform. However, within each stage of development, ranging from blister to dent, similar up-regulation in expression of many maize genes following inoculation with either method was observed; a total of 16 genes previously associated with resistance to pathogens were identified among the transcripts differentially expressed (DE) at p ≤ .05, FDR ≤ .10, and fold change ≥ 2.0 over all stages. The side needle technique produced a larger effect of infection as evidenced by 6,324 fungal reads versus 518 in silk channel and a higher level of aflatoxin. Correlations between approximately 7,000 fungal reads and the number of maize DE genes for each of the eight treatment groups was 0.56 (p = .152) and was 0.65 (p < .001) with levels of aflatoxin ranging from 0 to 137 ng g-1.

**Conclusion:** These correlations provided an internal measure of effectiveness of inoculation methods that were associated with the mostly up-regulation of defense-related genes in response to the presence of *Aspergillus flavus* in a unique maize genetic background.

## Introduction

Evaluation of differential gene expression (DGE) in maize in response to inoculation with *A. flavus* is challenging given the complexities of the pathogenic response under different environmental conditions and with the timing of infection and harvest. Studies up to the present, based on microarray analysis and/or RT-qPCR, have detected similar patterns of expression for some protein coding genes, such as chitinase, but many other detected genes have been novel to specific experiments. Since RNA-Seq can be applied without a reference genome, new gene sequences and sequence variations in the transcribed regions can now be recognized. In the case of maize, a reference genome is available (Cordes *et al.*, 2009) but given the polymorphism of the maize genome, arising from small and large-scale rearrangements (2-4), genes responsible for phenotypes of interest may not be present in the reference genome. TX772, a temperate Argentinian inbred featuring a hard and vitreous endosperm (Betrán, Isakeit and Odvody, 2002; Llorente *et al.*, 2010) does not share a common pedigree with most U.S. lines of Reid Yellow Dent, Lancaster or Iodent types (Goodman, 2005), and is genetically distant based on existing marker data (Smith, Murray and Heffner, 2015). Hybrids of Tx772 with BSSS germplasm (Stiff Stalk Synthetic) have exhibited high yield potential under irrigation. Most significantly for this study, Tx772 has shown good general combining ability to resist aflatoxin accumulation under non-wounding silk-channel inoculation, as evaluated using a diallel with certain tropical and subtropical inbred lines in several southern environments (Betrán, Isakeit and Odvody, 2002). In the multi-environment Southeast Regional Aflatoxin Trials (SERAT), the four hybrids using Tx772 as a parent ranked lower in aflatoxin contamination than 75-80% of the other test hybrids and ranged from 83-93% of the average check level (Wahl *et al.*, 2017).

To date, few studies have published profiles and changing patterns of differential gene expression in *developing* maize kernels (Lee *et al.*, 2002; Liu *et al.*, 2012; Lu *et al.*, 2013), and only one has reported on changes in expression following *A. flavus* infection (Dolezal *et al.*, 2014). From these studies, kernel maturity has been shown to be significantly associated with gene expression. Maize kernels go through stages of development beginning with silking (R1), followed by blister (R2), milk (R3), dough (R4), dent (R5) and physiological maturity (R6) (Ritchie, Hanway and Benson, 1997). In addition to the length of time following inoculation as a factor in percentage of kernels colonized and infected (Payne and Widstrom, 2008), changes in kernel biochemical composition during kernel development would be expected to influence the degree of infection by the fungus, and possibly the levels of aflatoxin contamination. For example, as early as 10 days after pollination (DAP) at the blister stage, certain alpha zeins and gamma zeins are only just detectable (Woo *et al.*, 2001) and reach significant levels by 15 DAP; in mature maize kernels at 55 – 65 DAP, prolamins or zeins comprise about 50% of the proteins (Liu *et al.*, 2008). Starch reserves are also built up in the endosperm from glucosyl, and fatty acids are stored in lipid bodies. This growing supply of nutrients can support the growth of the fungus which can begin with silk colonization (Payne and Widstrom, 2008) and has been found capable of infecting all tissues of immature kernels at different stages within 96 hours following infection (Dolezal *et al.*, 2013). (Dolezal *et al.*, 2014) further investigated this effect in a microarray analysis of transcriptional and physical changes in developing maize kernels infected by *A. flavus*, by inoculating ears of the inbred maize genotype B73 that is highly susceptible to the fungus and production of aflatoxin (Warburton *et al.*, 2013) at stages R2 – R5, and hand harvesting the ears four days later, followed by immediate RNA extraction. To gain a full spectrum of DGE relative to those mock-inoculated, Dolezal *et al.*, (2014) pooled all samples. Given the complexity of mechanisms occurring in three tissues: maternal, endosperm and embryo, this approach provided a more complete picture of all the genes likely to respond to the presence of the fungus at different stages in the field environment, although the tissues were not tested separately.

A different approach was taken in profiling gene expression with microarray analysis in *A. flavus* while colonizing maize kernels. Kernels were harvested from the blister to dent stages in the field, and then inoculated *in vitro* with fungal conidia (Reese *et al.*, 2011). Among the 190 fungal genes analyzed for patterns of expression, many exhibited differential expression (DE) that appeared to be dependent on the stage of maturity in the kernels.

Several methods have been developed and applied over the past forty years to inoculate maize through introducing *A. flavus* conidial spores in a suspension of distilled water to the ears of maize. Under natural conditions inoculation is otherwise governed strongly by the environment and presence and prevalence of fungal spores. Inoculation methods can be broadly grouped as “wounding” and “non-wounding”. Wounding methods include the side needle technique which involves inserting a needle under the husks and injecting about three ml. of suspension over the kernels, nicking some kernels in the process (Buckley, Williams and Windham, 2006), as well as pinbar inoculation and the knife technique (Scott *et al.*, 1991) both of which try to carry spores while creating a wound. A non-wounding technique traditionally used in Texas requires squirting the inoculum down the silk channel under the husk at the tip of the ear without nicking any kernels (Zummo and Scott, 1989), this is how Tx772 was identified as having decreased accumulation of aflatoxin. Another, more recent, non-wounding method is ground kernel inoculation which increases disease pressure by applying colonized kernels in the furrows to sporulate (Odvody *et al.*, 2000; Farfan *et al.*, 2015); this method is analogous to those used in current atoxigenic biocontrol applications of *A. flavus* (Isakeit *et al.*, 2010; Grubisha and Cotty, 2015). Non-wounding methods are likely better choices for grower-relevant testing of maize in preharvest aflatoxin susceptible production areas where populations of wounding insects are low and wind or other non-wounding natural inoculation are more important. In contrast, the wounding technique typically can produce higher levels of aflatoxin (Buckley, Williams and Windham, 2006).

The primary objective of this exploratory study was to identify genes significantly differentially expressed between inoculated and non-inoculated Tx772 kernels at a given stage of maturity (blister, milk, dough or dent), focusing on validating those genes previously reported as having contributed to a pathogenic response. A second objective was to compare the effects of two inoculation methods and the non-inoculated control on gene expression at a given stage of maturity, with respect to identity, function and degree of fold change, as well as to determine if one inoculation method resulted in a larger number of consistently significantly differentially expressed genes (DEGs). In conjunction with this assessment, levels of aflatoxin, and the variation and characterization of fungal transcripts in each sample were also determined for evaluation of inoculation success.

## Methods and Materials

### Inoculum preparation and application

Inoculum was prepared from the *A. flavus* isolate NRRL 3357 grown on sterilized corn kernels. The conidia were washed off and purified by repeated sedimentation through centrifugation at 4°C to obtain a final spore concentration of 10^7^mL^−1^(Wahl *et al.*, 2017). The same inoculum was used for silk channel and side needle inoculation.

Replicate samples of maize inbred line TX772 (Llorente *et al.*, 2010) were grown in College Station in 2012 and subjected to one of three treatments: 1. Three ml of inoculum down the silk channel (Zummo and Scott, 1989). Three ml of inoculum by side needle (Buckley, Williams and Windham, 2006), or 3. no inoculation at 10 days after pollination (DAP). Ears were harvested at the following stages of maturity: 1. blister (11dap) 2. milk/early dough (21-25 dap) 3. late dough (29 dap) and 4. early dent (36 dap). Since the milk and early dough stages were only four days apart, these samples were analyzed as one group named “milk”. All kernels cut from each ear were flash frozen at harvest at −80C, ground with a mortar and pestle, and thoroughly mixed for RNA extraction and testing for the levels of aflatoxin.

### RNA extraction and sequencing

Total RNA was extracted from 40 mg of finely ground kernel samples using the Spectrum™ Plant Total RNA Kit; Sigma-Aldrich, St. Louis, MO, 2010, according to manufacturer’s protocol., except for some samples in lysis solution that needed to be filtered twice. The total RNA was qualified and quantified with an Experion RNA HighSens Analysis Kit; Bio-Rad, Hercules, CA. Approximately 1.4 µg of total RNA was used for cDNA synthesis, followed by construction of RNA-Seq libraries using the TruSeq RNA kit version 2.0, Illumina (San Diego, CA). Samples were submitted for sequencing at BGI Americas (Cambridge, MA) using a module of 100PE (paired ends) on the Illumina HiSeq 2000 platform. The clean reads were sorted according to the barcode of its library and extracted using the BGI pipeline. These clean reads were deposited at https://www.ncbi.nlm.nih.gov/sra, with Project Number PRJNA384648.

### Transcriptome assembly and quality assessment

A *de novo* assembly of transcripts and genes were made at the High Performance Research Computing resources at Texas A&M University through application of the Trinity platform (Haas *et al.*, 2013) on the 24 samples to form the basis of a single Trinity.fasta file. Twenty-four RSEM gene and isoform results files were generated from the Trinity.fasta file in conjunction with the 48 left and right compressed fastq files. Trinity functions were run to compare biological replicates and determine the relatedness of samples. A decision was made to eliminate one silk channel sample and one non-inoculated sample from the milk group in the differential expression analysis that were clear outliers as shown in the principal component analysis (Figure 1) a. The blister stage consisted only of the three treatments without replicates, there were three replicates at the milk stage, and two replicates each at the dough and dent stages.

**Fig 1.**
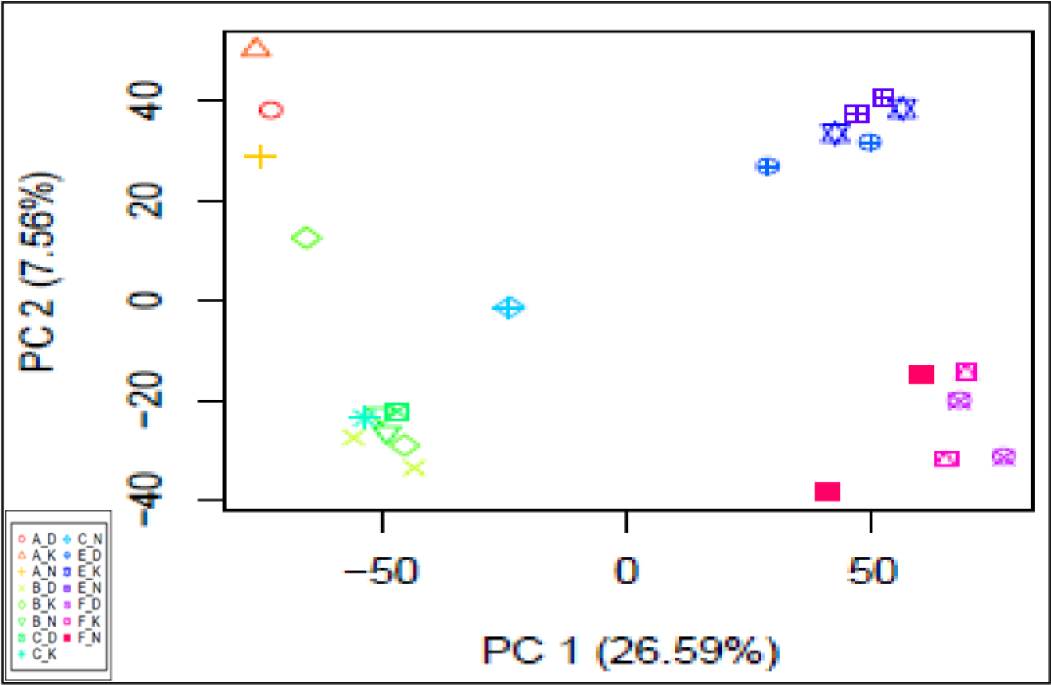
PCA of differential gene expression of maize kernels under different treatments. First letter refers to harvest dates, second letter refers to inoculation treatments of kernels. A-Jun 08, B&C-Jun 18 & 22, E-Jun 26, F - Jul 3, D-side needle, K-silk channel.

Because the mRNA from a fungus-inoculated sample was a mixture of mRNA’s from the host plant and pathogen, fungal sequences were identified in the Trinity.fasta file through the application of BLAST+ against the downloaded cDNA file of *Aspergillus flavus* (NRRL3357) which was obtained from the website: http://fungi.ensembl.org/info/website/ftp/index.html. A maximum p-value for identifying *A. flavus* cDNA sequences was set to 1e −20. These fungal sequences were characterized in BLASTn and summarized in S2 Table, and their Trinity ids were used as a filter to remove them from the maize count matrices.

### Statistical models and differential gene expression

Differential gene expression analysis was conducted using edgeR from Bioconductor (Haas *et al.*, 2013) and run independently of Trinity for greater flexibility in statistical modeling. This R package references a table of actual (or expected) read counts with columns corresponding to the sample libraries, and rows corresponding to the assembled transcripts. The application of a negative binomial distribution in this package assumes that the true gene abundances follow a gamma distribution across replicate samples. A series of functions in R were designed to call upon edgeR routines to: 1) filter out lowly expressed genes with read counts less than 5; 2) implement the experimental design which consisted of two main types of contrasts for differential gene expression: one maturity level versus another, and one inoculation method versus non-inoculated, at each maturity level; 3) calculate and apply trimmed mean of M-values (TMM) normalization scaling factors to correct for the differences in library sizes (Dillies *et al.*, 2013); 4) estimate gene-specific dispersion appropriate for the negative binomial model to account for the biological coefficients of variation expected to exist among genes; 5) conduct a likelihood ratio test (LRT) to determine significant differential gene expression defined by p≤.05, FDR ≤ .10 and log2FC ≥ 2; and finally 6) Run a modified “ TopTags” in edgeR to find the “n” most significant DEGs, in which the Benjamin-Hochberg (BH) method is applied to control the false discovery rate (Benjamini and Hochberg, 2018). In 4), the gene-specific dispersion factor is calculated on the entire set of samples, at first as a common factor to all genes in all samples, followed by a tagwise dispersion. This provides some correction to gene expression measured for treatments that are lacking replicates, as was the case with the blister group.

A paired t-test was applied to average numbers of reads among replicates for each gene between samples inoculated by silk channel with those inoculated by side needle at each level of maturity, using a similar statement in R: t.test(blist_side, blist_silk, paired = TRUE).

### Identification of differentially expressed genes

The primary database referenced for identification of each significant differentially expressed gene as represented by a Trinity.fasta sequence was MaizeGDB (Lawrence *et al.*, 2008) that provided Gramene numbers based on v4 of the maize B73 reference genome. The database most commonly accessed within MaizeGDB was MaizeCyc (Monaco *et al.*, 2013), that provided the name of the most likely gene product, but often Pfam (Finn *et al.*, 2014) and InterPro (Hunter, 2002; Apweiler *et al.*, 2014) were checked as well. NCBI BLASTn (Johnson *et al.*, 2008) was consulted for sequences with alternative characterizations. Original articles were also referenced for gene identities, function and biological processes, especially with respect to pathogenic responses and disease resistance.

### Measurement of aflatoxin contamination

Sub-samples of kernels ground for each treatment ranging from 16 to 35 g per ear were tested for aflatoxin concentration using the VICAM AflaTest^®^per manufacturer’s instructions and as used and described in more detail in (Wahl *et al.*, 2017).

## Results and Discussion

### Statistics on RNA-Seq transcript and gene assembly

A total of 313.8 million clean reads were obtained from the sequencing of the 24-RNA-Seq libraries constructed from 24 samples. The clean reads had a quality of > Q20 for an average of 96.64% of the reads ranging from 95.95 - 97.35%. Each sample had an average of 13.1 million clean reads, varying from 10.6 – 13.7 million. A total of 268,720 transcripts (isoforms) resulting from the transcriptome assembly were associated with 152,574 “genes” with an average contig length of 636 bases. Eighty-five percent of the reads aligned concordantly more than one time, and twelve percent aligned exactly once.

The number of paired-end reads with at least some overlap counted by edgeR in each sample (Table 1 a and b), averaged about 5.6M, and the number of transcripts or loci identified by *de novo* assembly in Trinity ranged from 57.8K – 77.8K over all samples. The total number of DEGs under silk channel (non-wounding) inoculation was 315, while side needle (wounding) was 457 with the breakdown according to development stage displayed in Table 1 (d).

**Table 1.** Numbers of reads and transcripts by sample

| Sample | Read counts (a) | Number of transcripts (b) | Maize DEGs per treatment (c) † | Number of fungal reads (d) | Total reads per type of inoculation (e) | Aflatoxin ng g-1 (f) |
| --- | --- | --- | --- | --- | --- | --- |
| AD_1 | 5,831,702 | 62,308 | 70 | 1,924 | 1,924 | 137.7 |
| AK_1 | 5,795,934 | 60,048 | 261 | 14 | 14 | N/A |
| AN_1 | 5,899,959 | 59,480 |  | 0 |  | 0 |
| BD_1 | 5,904,363 | 64,535 |  | 305 |  | 27.1 |
| BD_2 | 5,793,513 | 63,711 |  | 49 |  | 0.0 |
| BD_3 | 5,887,542 | 61,770 | 18 | 167 | 521 | 7.8 |
| BK_1 | 5,726,260 | 61,295 |  | 9 |  | 6.1 |
| BK_2 | 4,706,782 | 57,752 |  | 13 |  | 0.0 |
| BK_3 | 5,895,009 | 59,826 | 3 | 6 | 28 | 3.0 |
| BN_1 | 5,009,778 | 62,057 |  | 5 |  | 0 |
| BN_2 | 5,181,385 | 58,756 |  | 12 |  | 0 |
| BN_3 | 5,718,067 | 63,218 |  | 1 |  | 0 |
| ED_1 | 5,881,636 | 71,732 |  | 715 |  | 8.9 |
| ED_2 | 5,932,908 | 68,566 | 16 | 240 | 955 | 0.0 |
| EK_1 | 5,450,481 | 67,248 |  | 187 |  | 2.7 |
| EK_2 | 4,943,068 | 62,714 | 20 | 107 | 294 | 1.8 |
| EN_1 | 5,302,995 | 66,341 |  | 3 |  | 0 |
| EN_2 | 5,779,269 | 64,644 |  | 8 |  | 0 |
| FD_1 | 5,855,546 | 80,208 |  | 2,704 |  | 21.7 |
| FD_2 | 5,894,548 | 75,818 | 353 | 220 | 2924 | 3.7 |
| FK_1 | 5,849,170 | 77,779 |  | 93 |  | 10.8 |
| FK_2 | 6,069,033 | 76,825 | 31 | 89 | 182 | 3.2 |
| FN_1 | 6,002,760 | 74,868 |  | 28 |  | 0 |
| FN_2 | 5,966,850 | 77,514 |  | 30 |  | 0 |

† Differentially expressed genes at $p \leq .05$ , and $FDR \leq .10$ , and $\text{Log}_2(FC) \geq 1.0$
Stages of seed development: A - blister B - milk, E - dough, F - dent
Method of inoculation: D - side needle, K - silk channel
(a) Read counts reflect number of *pairs* of reads that overlap a fragment of cDNA.

### General characteristics of gene expression

The maturity of the kernels at harvest appeared to be the greatest source of variation, as indicated by principal component analysis of log2(read counts), re-scaled or normalized based on library sizes in Figure 1. Using a single gene as an example, at the sucrose synthase (shrunken1) locus, the read counts normalized by the Trimmed Mean of *M*-values (TMM) method (Dillies *et al.*, 2013) at the blister stage ranged from 1,032 - 1,217, at the milk stage: 444 - 701, at the dough stage: 195 - 276, and at the dent stage: 161 – 259. This led samples to primarily be grouped and analyzed according to the stage of development, which indicated effects of any other treatments or factors should be evaluated within the context of stage of maturity.

Even with the limited number of biological replicates within treatment sets, the DGE in response to inoculation and/or the presence of *A. flavus* were distinctive at each stage regarding specific patterns of gene ontologies, numbers of genes, and magnitude and direction of expression. A paired t-test applied to the average read counts at each stage showed no significant differences between the two inoculation methods for blister and dent samples. However, results were significant at the milk and dough stages, likely due to fewer genes being differentially expressed, and of these, only a small percentage were up or down-regulated under both types of treatment. In the blister group, both methods of inoculation versus non-inoculation resulted in mostly up-regulation of over 50 genes, with similar fold changes between inoculation methods that often exceeded two; this despite the observation that few fungal reads were detected in the silk channel sample. This finding suggests the fungus can dramatically influence the host’s gene expression, even at low levels. More noteworthy is the DGE in response to the inoculation at such an early stage of development, as fungal infection of kernels damaged (Dolezal *et al.*, 2014) or un-damaged (Marsh and Payne, 1984) have not been reported and have not been believed to occur before milk stage. However, it is possible that early host responses to inoculum is a function of a “resistant” genotype that would not be observed in a susceptible genotype.

In total, there were 685 unique transcripts DE between inoculated samples versus non-inoculated samples over all stages of maturity within the bounds of p ≤ .05, FDR ≤ .10, and a fold change of 2 (log2FC ≥ 1). The likelihood ratio test was applied to each comparison which permitted a ranking of differentially expressed genes (DEGs) and provided a p-value and FDR. In Table 2, the fifty genes most significantly differentially expressed at the blister stage under both methods of inoculation are presented, excluding those with uncharacterized gene products.

**Table 2.**
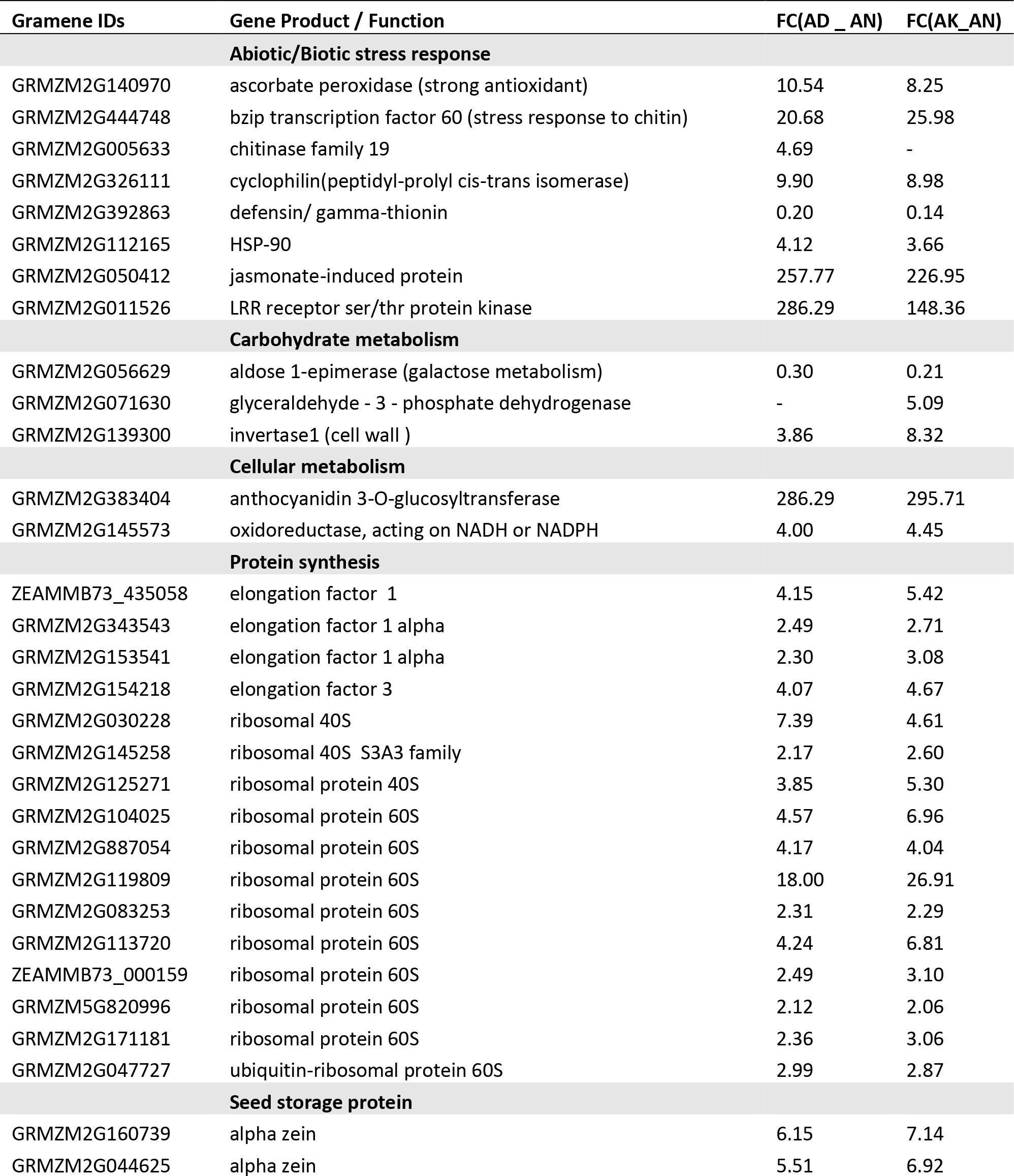
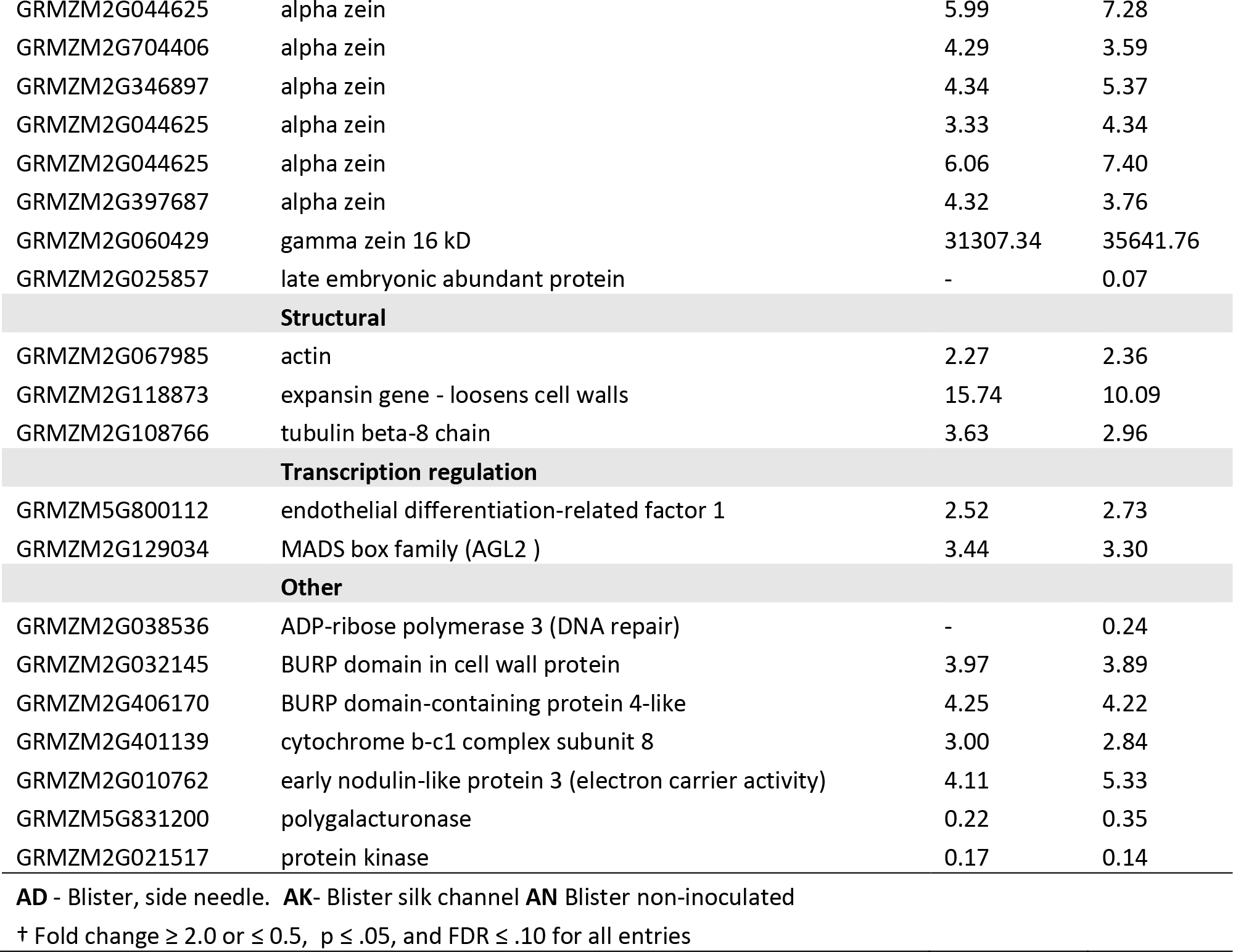
Differential gene expression (Fold Change †) in the silk channel or side needle inoculated kernels vs non-inoculated kernels at the blister stage

Forty-four genes were significantly DE under *both* inoculation treatments, with similar fold changes and direction. In Tables 3 and 4, DGE of kernels at stages milk to dent are represented in a similar manner; the largest group of significantly DEGs was found in the side-needle inoculated dent kernels, but no more than the top fifty genes were subjected to analysis. Graphical depiction of numbers of up-regulated and down-regulated genes in Tables 2-4 are shown in Figures 2a and 2b.

**Table 3.**
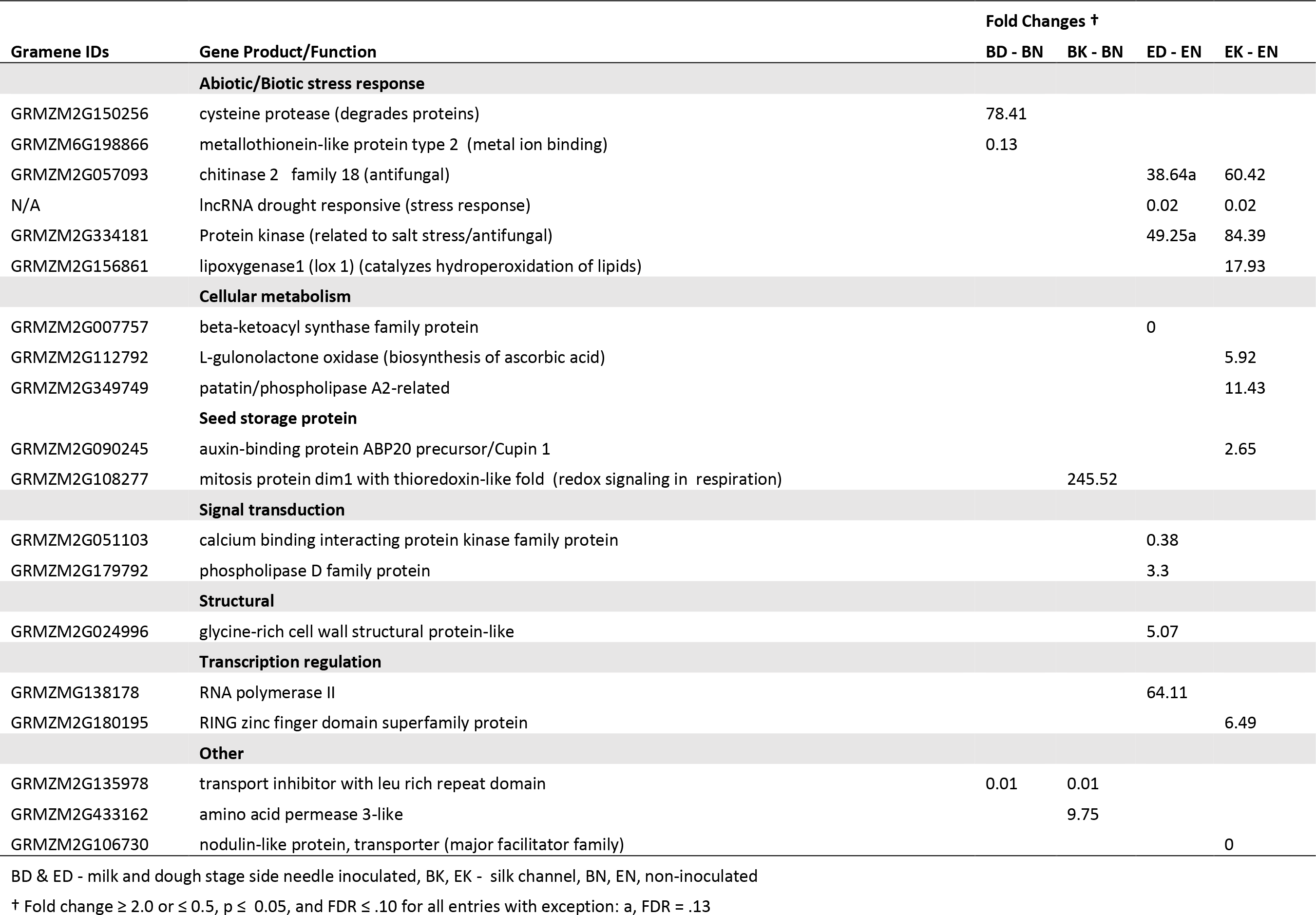
Differential gene expression (fold change †) in silk channel or side needle inoculated kernels vs non-inoculated kernels at milk and dough stage

| Gramene IDs | Gene Product/Function | Fold Changes † |  |  |  |
| --- | --- | --- | --- | --- | --- |
|  |  | BD - BN | BK - BN | ED - EN | EK - EN |
|  | Abiotic/Biotic stress response |  |  |  |  |
| GRMZM2G150256 | cysteine protease (degrades proteins) | 78.41 |  |  |  |
| GRMZM6G198866 | metallothionein-like protein type 2 (metal ion binding) | 0.13 |  |  |  |
| GRMZM2G057093 | chitinase 2 family 18 (antifungal) |  |  | 38.64a | 60.42 |
| N/A | lncRNA drought responsive (stress response) |  |  | 0.02 | 0.02 |
| GRMZM2G334181 | Protein kinase (related to salt stress/antifungal) |  |  | 49.25a | 84.39 |
| GRMZM2G156861 | lipoxygenase1 (lox 1) (catalyzes hydroperoxidation of lipids) |  |  |  | 17.93 |
|  | Cellular metabolism |  |  |  |  |
| GRMZM2G007757 | beta-ketoacyl synthase family protein |  |  | 0 |  |
| GRMZM2G112792 | L-gulonolactone oxidase (biosynthesis of ascorbic acid) |  |  |  | 5.92 |
| GRMZM2G349749 | patatin/phospholipase A2-related |  |  |  | 11.43 |
|  | Seed storage protein |  |  |  |  |
| GRMZM2G090245 | auxin-binding protein ABP20 precursor/Cupin 1 |  |  |  | 2.65 |
| GRMZM2G108277 | mitosis protein dim1 with thioredoxin-like fold (redox signaling in respiration) |  | 245.52 |  |  |
|  | Signal transduction |  |  |  |  |
| GRMZM2G051103 | calcium binding interacting protein kinase family protein |  |  | 0.38 |  |
| GRMZM2G179792 | phospholipase D family protein |  |  | 3.3 |  |
|  | Structural |  |  |  |  |
| GRMZM2G024996 | glycine-rich cell wall structural protein-like |  |  | 5.07 |  |
|  | Transcription regulation |  |  |  |  |
| GRMZMG138178 | RNA polymerase II |  |  | 64.11 |  |
| GRMZM2G180195 | RING zinc finger domain superfamily protein |  |  |  | 6.49 |
|  | Other |  |  |  |  |
| GRMZM2G135978 | transport inhibitor with leu rich repeat domain | 0.01 | 0.01 |  |  |
| GRMZM2G433162 | amino acid permease 3-like |  | 9.75 |  |  |
| GRMZM2G106730 | nodulin-like protein, transporter (major facilitator family) |  |  |  | 0 |

BD & ED - milk and dough stage side needle inoculated, BK, EK - silk channel, BN, EN, non-inoculated
† Fold change $\geq 2.0$ or $\leq 0.5$ , $p \leq 0.05$ , and $FDR \leq .10$ for all entries with exception: a, $FDR = .13$

**Table 4.** Differential gene expression (fold change †) in silk channel or side needle inoculated kernels vs non-inoculated kernels at dent stage

|  |  | Fold Changes † |  |
| --- | --- | --- | --- |
| Gramene IDs | Gene Product/Function | FD - FN | FK - FN |
| Abiotic/Biotic stress |  |  |  |
| GRMZM2G172204 | beta glucosidase jasmonate-induced aggregating factor1 | 8.87 | 5.21b |
| GRMZM2G042639 | GST activity safener induced1 | 2.19 | 1.98 |
| GRMZM2G127251 | hydroxycinnamoyl transferase3 | 3.28 |  |
| GRMZM5G822593 | lipoxygenase 8, PLAT domain | 6.47 | 3.51b |
| GRMZM2G152638 | lncRNA (drought responsive) | 3.68 | 2.28 |
| GRMZM2G036048 | O-methyltransferase (lignin biosynthesis) | 5.4 |  |
| GRMZM2G127418 | O-methyltransferase (lignin biosynthesis) | 6.81 | 3.75b |
| Carbohydrate metabolism |  |  |  |
| GRMZM2G103055 | alpha amylase | 3.82 |  |
| GRMZM2G138468 | alpha amylase3 | 48.1 | 34.02 |
| GRMZM2G031660 | beta-glucosidase | 0.33 |  |
| GRMZM2G410916 | glycosyl transferase | 0.42 |  |
| GRMZM2G394450 | invertase (Ivr1) | 72.97 |  |
| GRMZM2G394450 | invertase (Ivr1) | 5.96 |  |
| GRMZM2G336448 | carbohydrate transporter | 1.82 |  |
| GRMZM2G383404 | anthocyanidin 3-O-glucosyltransferase |  | 0.09 |
| GRMZM2G168474 | O-glucosyltransferase 2 (cis-zeatin ) |  | 2.02 |
| Other cellular metabolism |  |  |  |
| GRMZM2G112792 | L-gulonolactone oxidase (biosynthesis of ascorbic acid) | 11.35 | 5.79 |
| GRMZM2G349749 | patatin/phospholipase A2-related | 5.03 |  |
| GRMZM2G153536 | amino-acid aminotransferase (branched-chain) | 2.47 |  |
| GRMZM2G359298 | copper amine oxidase | 0.41 |  |
| GRMZM2G170400 | cyclopropane-fatty-acyl-phospholipid synthase | 6.32 |  |
| GRMZM2G125268 | aldehyde dehydrogenase2 | 9.28 |  |
| GRMZM2G023847 | blue copper protein (redox process in photosynthesis) | 6.11 |  |
| GRMZM2G077809 | copine (Ca2+ dependent phospholipid-binding) | 0.01 |  |
| GRMZM2G332562 | dicarboxylic acid transport (transports amino acids) | 2.22 |  |
| GRMZM2G076239 | hydroxyacid oxidase 1 (glycolate oxidase) | 0.3 |  |
| GRMZM5G842071 | laccase 1 (LAC1) gene, a multicopper oxidase | 2.69 |  |
| GRMZM5G889138 | NADH dehydrogenase subunit 7 | 5.27 |  |
| Protein degradation |  |  |  |
| GRMZM2G029506 | peptidase activity ATP-dependent | 1.98 |  |
| GRMZM2G124684 | proteinase - Aspartic | 506.74 |  |
| GRMZM2G164787 | polyubiquitin containing 7 ubiquitin monomers |  | 69.17 |
| GRMZM2G016323 | ubiquitin carboxyl-terminal hydrolase 23 |  | 0.02 |
| GRMZM2G075104 | ubiquitin conjugation factor E4 |  | 49.86 |
| GRMZM2G037411 | pectinesterase | 0.38 |  |
| **Protein synthesis** |  |  |  |
| GRMZM2G474534 | ribosomal protein 30S , chloroplastic | 46.85 |  |
| GRMZM2G000741 | GTPase Protein-synthesizing |  | 0.01 |
| **Seed storage protein** |  |  |  |
| GRMZM2G090245 | auxin-binding protein ABP20 precursor/Cupin 1 | 3.44 |  |
| GRMZM2G044625 | alpha zein | 0.37 |  |
| GRMZM2G044625 | alpha zein | 0.33 |  |
| GRMZM2G397687 | alpha zein | 0.4 |  |
| ZEAMMB73_324839 | cupin | 17.72 |  |
| GRMZM2G060429 | gamma zein 16 kDa | 0.25 |  |
| GRMZM2G174883 | legumin 1,cupin 1 domain | 0.43 |  |
| **Signal transduction** |  |  |  |
| GRMZM2G122228 | protein phosphatase homolog8 | 0.36 |  |
| GRMZM2G025459 | protein kinase 5'-AMP-activated subunit beta-1 | 2.95 |  |
| **Transcription regulation** |  |  |  |
| GRMZM2G180195 | RING zinc finger domain superfamily protein |  |  |
| GRMZM5G822829 | anthocyanin pathway r1 (colored1) |  | 0.47 |
| GRMZM2G146283 | endosperm-specific prolamin box binding factor (PBF) zinc finger | 0.4 | 0.46 |
| GRMZM2G006676 | ligand-dependent nuclear receptor activity | 2.27 |  |
| GRMZM2G359952 | MADS2 | 0.3 |  |
| GRMZM2G172327 | MYB-transcription factor 14 | 0.36 |  |
| GRMZM2G015534 | opaque endosperm2 Basic leucine-zipper C terminal | 0.49 |  |
| GRMZM2G157219 | trihelix-transcription factor 1 | 0.01 |  |
| **Other** |  |  |  |
| GRMZM2G106730 | nodulin-like protein, transporter (major facilitator family) |  |  |
| GRMZM2G061527 | leucine-rich repeat domain | 3.18 |  |
| GRMZM2G073114 | ripening-related protein 3-like | 134.21 |  |
| GRMZM2G430755 | cation/H(+) antiporter 15-like |  | 2.18 |
| GRMZM2G076343 | legume lectins beta domain containing protein |  | 2.42 |
| GRMZM2G017013 | protein binding (in apoptosis) |  | 0.18 |

FD - dent side needle inoculated, FK - silk channel, FN - non-inoculated
† Fold change $\geq 2.0$ or $\leq 0.5$ , $p \leq 0.05$ , and $FDR \leq .10$ for all entries with exceptions listed below:
a $FDR = .13$ , b $.13 \leq FDR \leq .20$

**Fig 2a and 2b.**
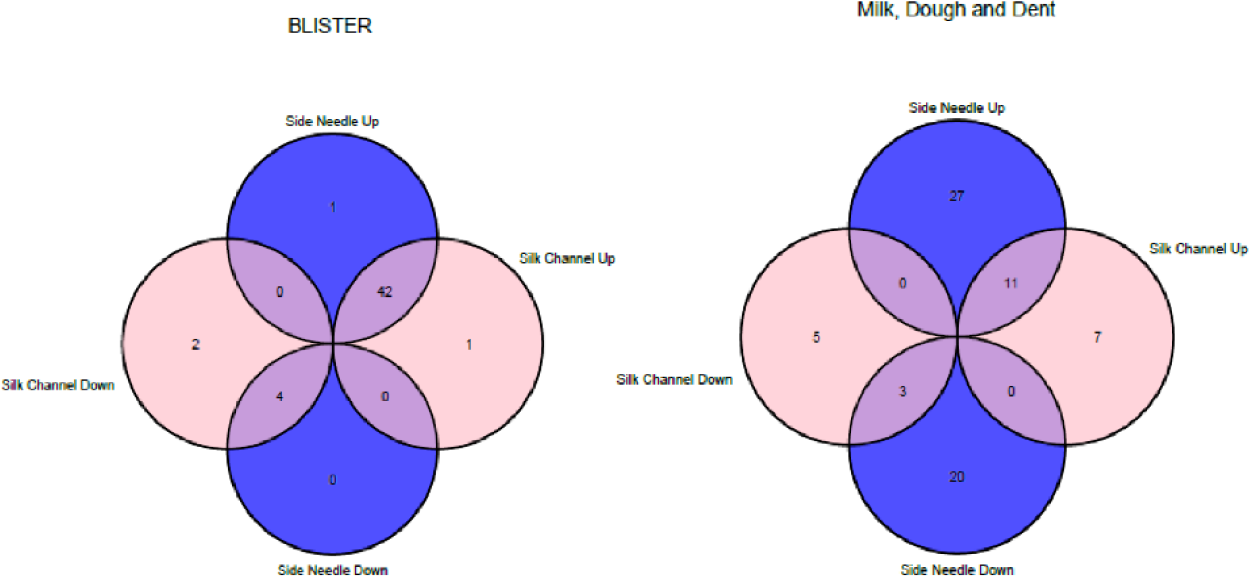
Comparison of inoculation methods on numbers of up-regulated or down-regulated genes relative to the non-inoculated samples as shown in Tables 2 and 3. (Venn Diagram designs and code provided by Lambirth KC, Whaley AM, Blakely IC, Schleuter JA, Bosst KL, Loraine AE and Piller KJ. A comparison of transgenic and wild soybean seeds: analysis of transcriptome profiles using RNA-Seq. BMC Technology. 2015; 15(80): 1-17).

Over all stages of development, aflatoxin levels were higher in the side needle inoculated samples, and there were much higher levels of fungal reads in certain side needle samples.

### DEGs previously associated with injury or pathogenesis and their function in A. flavus infection

A total of 16 DE protein-coding genes were identified under one or both inoculation treatments in this study, each of which has been previously associated with a response to injury and/or presence of a pathogen in relevant other studies (S1 Table). These primarily increased under inoculation but one, gamma-thionin, decreased (Figure 3). Given the corroborating evidence of our study, these 16 DE protein-coding genes are worth discussing in more detail and will be grouped according to a function highlighted in this study.

**Fig 3.**
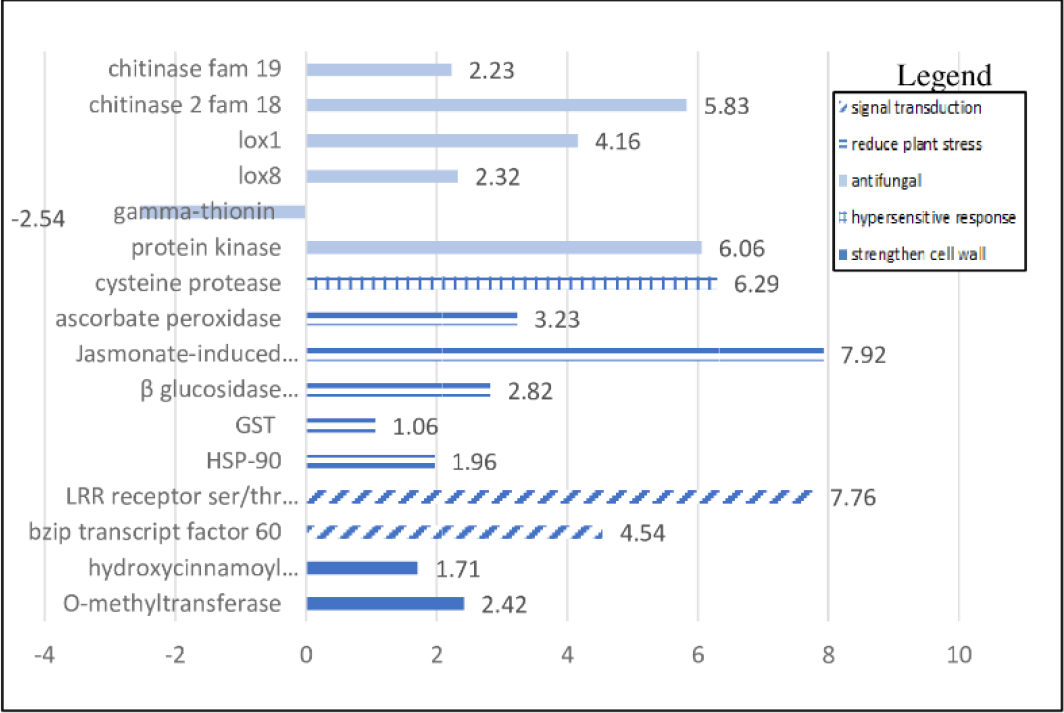
Differential gene expression represented by log2 fold changes at different stages of kernel maturity that have been previously associated with a biotic stress response, at p ≤ .05, and FDR ≤ .10.

#### Antifungal group

Chitinases are often implicated in plant fungal defenses and belong to the second largest group of pathogenesis-related proteins (PR). They are included in families 18 and 19 of glycoside hydrolases and catalyze chitin degradation in the fungal cell wall (Ferreira *et al.*, 2007). Some classes are mainly chitin-binding, and thus inhibit fungal growth by disrupting cell polarity when bound to the fungal cell wall. In this study, GRMZM2G005633 of family 19 was up-regulated by over four-fold in the side-needle inoculated kernels at blister stage, while in the dough stage both the silk channel and side needle inoculations were associated with up-regulation of about 32-fold in GRMZM2G057093 of family 18. The former was assigned to the maize genome region of bin 10.04 (Hawkins et al, 2015) while the latter was assigned to bin 1.08 by MaizeCyc, a network of metabolic pathways delineated in B73 (Monaco *et al.*, 2013). Hawkins *et al.*, (2015) did not find that chitinase (GRMZM2G005633), which they identified in their QTL mapping populations of hybrids derived from susceptible and resistant parents, contributed to any phenotypic effect with respect to aflatoxin contamination resistance. This was possibly due to post-translational modifications of the chitinase by a fungal protease (Naumann, Wicklow and Kendra, 2009; Naumann and Wicklow, 2010). Chitinase 2 (GRMZM2G057093), significantly DE in this study, did not appear in the genetic mapping populations discussed previously, likely because it was not segregating between the parents (Hawkins *et al.*, 2015). This gene was expressed in the dough samples and may have been instrumental in preventing the levels of infection from more quickly reaching those of the blister or dent stages, according to the relative numbers of fungal reads as will be discussed later.

Resistance to pathogens exhibited by most lipoxygenases have often been difficult to determine (Fountain *et al.*, 2014), including the two up-regulated in this study, LOX1 (GRMZM2G156861) under silk channel inoculation in dough kernels by 16-fold, and LOX8 (GRMZM5G822593) by more than four-fold in the dent samples. In contrast to our results, a different maize line, Mp719, also known to be relatively aflatoxin resistant revealed a downregulation of expression of LOX1 upon inoculation with the fungus relative to mock-inoculated with water, while LOX8 was significantly up-regulated (Ogunola *et al.*, 2017). There is evidence also from a QTL study (Mideros *et al.*, 2014) that LOX8 contributes to resistance to aflatoxin to resistance to aflatoxin contamination. Furthermore, in a genome-wide association study to identify metabolic pathways contributing resistance to aflatoxin contamination in maize, genes for both LOX1 and LOX8, contributed a highly significant positive effect (Tang *et al.*, 2015). LOX8, among other lipoxygenases, contributes to the biosynthesis of the hormone jasmonate (JA) (Christensen *et al.*, 2013), and this is the key hormone associated with an incremental decrease to levels of aflatoxin in a GWAS panel (Tang *et al.*, 2015).

One protein belonging to a multi-functional class of defense proteins, flower-specific γ-thionines (GRMZM2G392863), was down-regulated in the inoculated blister samples. This was the only DE protein identified as antifungal and protective against insect pests to be downregulated under inoculation (Lay *et al.*, 2003) in this study. Another unnamed protein, (GRMZM2G334181) related to a protein kinase that responds to salt stress (Zhang *et al.*, 2009) shows homology to an antifungal protein with protease inhibitory activity (Sawano *et al.*, 2007) as aligned in the Pfam database, was similarly DE in the dough stage under both treatments by around six-fold.

#### Hypersensitive type

There are many normal metabolic processes that can lead to the production of reactive oxygen species (ROS) that are harmful to the cell, but abiotic and biotic stresses may lead to an excess of ROS (Shigeoka *et al.*, 2002). If ROS levels exceed a certain threshold programmed cell death (PCD) or apoptosis can be activated in a hypersensitive response to an invading pathogen (Solomon *et al.*, 1999). Cysteine protease (GRMZM2G150256), which is activated by high levels of ROS, was highly up-regulated in side needle inoculated milk stage samples. Plants also have protease inhibitors to limit the PCD, but none were DE in this study.

#### Stress response group

In contrast to the effects of proteases, peroxidases in the plant as well as in the fungus are protective against the damaging effects of ROS arising from increased levels of H^2^O^2^ in response to pathogenic attacks and other factors (Shigeoka *et al.*, 2002), and an ascorbate peroxidase (APX) gene (GRMZM2G140970) was up-regulated by about 8-fold in both blister samples. Experiments on transgenic antisense tobacco with reduced APX infected with the bacterium *Pseudomonas syringae* resulted in elevated cellular H^2^O^2^ levels that led to enhanced cell death (Mittler *et al.*, 1999). However, it must be considered that H^2^O^2^ has beneficial roles as well as detrimental ones that must be balanced in a regulatory system (Shigeoka *et al.*, 2002); this would explain why gene expression of some peroxidases are down-regulated instead of up-regulated under similar experimental conditions.

Jasmonate is a plant hormone noted for increasing resistance to necrotrophs such as *A. flavus*, as opposed to biotrophs that obtain nutrients from living tissue (Glazebrook, 2005). In this study two genes associated with proteins described as “induced by jasmonate” were up-regulated, one in the blister inoculated samples (GRMZM2G050412) with large positive fold changes for side needle and silk channel treatments, and the other a beta glucosidase jasmonate-induced aggregating factor1 (GRMZM2G172204) in the dent samples with smaller fold changes.

Glutathione-S-transferases (GST) has a role in detoxifying toxic substances encountered during biotic and abiotic stress and has been moderately correlated with resistance to a number of maize pathogens (Wisser *et al.*, 2011). Gene expression for a protein with GST activity (GRMZM2G042639) was upregulated about two-fold under both inoculation treatments in the dent samples.

Certain heat shock proteins such as Hsp90 are involved in disease and pest resistance besides acting as molecular chaperones to regulate and maintain proper protein conformations (Liu *et al.*, 2004; Xu *et al.*, 2012). Kelley *et al.*, (2012) reported that Hsp90 was up-regulated in resistant Mp313E over susceptible Va35 in response to *A. flavus* inoculation. In this study Hsp90 GRMZM2G112165) was differentially expressed in the inoculated blister samples about four-fold.

#### Signal transduction group

The complex roles of plant receptor-like serine threonine kinases with a leucine rich repeat domain have often been related to their activities in signaling and plant defense (DeYoung and Innes, 2006; Afzal, Wood and Lightfoot, 2008). One study demonstrated the contribution to resistance to powdery mildew caused by the fungus, *Blumeria graminearum* in wheat (Chen *et al.*, 2016), and another to rust resistance in barley stems (Brueggeman *et al.*, 2002). In the blister samples, the fold change for the LRR receptor ser/thr protein kinase (GRMZM2G011526) was an approximately 64-fold increase for both treatments.

Bzip transcription factor 60 (GRMZM2G444748), identified in *Arabidopsis thaliana* as AT1G42990 has shown significant DGE under pathways unique in the response to chitin elicitation (Libault *et al.*, 2007; Zhang *et al.*, 2007) that were initially independent of three stress hormones: ethylene, jasmonic acid and salicylic acid. Bzip transcription factor 60 was DE by more than 20-fold in both the silk and side needle blister samples (Table 2).

#### Lignin biosynthesis group

In host plants, a build-up of lignin has been associated with resistance to fungal growth, providing a first line of defense against pathogen invasion as it strengthens the cell wall against mechanical pressure arising from fungal appressoria attempting penetration. Lignin is synthesized from phenylpropanoid hydroxycinnamyl alcohols, (Ebrahim, Usha and Singh, 2011), and the enzyme hydroxycinnamoyl transferase (GRMZM2G127251) was up-regulated by more than two-fold in the dent side-needle inoculated samples. Another enzyme involved in the lignin biosynthesis pathway, O-methyltransferase (GRMZM2G127418, GRMZM2G036048), was up-regulated by more than four-fold in both side needle and silk channel samples at the dent stage. Regarding hydroxycinnamoyl transferase3 involved in lignin biosynthesis, (Kelley *et al.*, 2012) reported that a related gene, cinnamoyl-CoA reductase was significantly expressed in the susceptible maize line Va35 upon inoculation with *A. flavus*. O-methyltransferase is also essential in lignin biosynthesis, and a mutation in that gene produces the *brown midrib3* phenotype, which modifies and reduces lignin content in the stems and roots, making the stems more digestible as a forage crop (Vignols *et al.*, 2007). The effects of RNA-mediated silencing to inhibit the former enzyme, which has a key position in the phenylpropanoid pathway in the formation of lignin arrested early development of Arabidopsis plants (Hoffman et al., 2004). In transgenic lines of alfalfa (*Medicago sativa* L.) that limited enzyme activity from 15-50% resulted in significant stunting, reduced biomass, and delayed flowering (Shadle *et al.*, 2007). Therefore, knocking down enzymes that are known to be key to lignin biosynthesis to test disease resistance would likely have undesirable side effects. However, the correlation between levels of phenolic compounds that are incorporated into lignin following inoculation with the fungus as was done with peanuts, another crop subject to aflatoxin contamination, has been measured (Liang et al., 2006). Not only was there a significant negative correlation between infection rate and lignin abundance seven days following inoculation, but resistant genotypes required much less time to reach maximum levels of key enzyme activity to metabolize lignin precursors than susceptible types. Other studies have shown this as well (Fajardo *et al.*, 1995, 2009; Liang, Luo and Guo, 2006).

### Other classes of genes DE not directly related to pathogenic response

In addition to the proteins described above for which there is some evidence of an antifungal effect, two other classes of genes that were differentially expressed in this and previous studies are likely important.

Five different alpha zein genes in the blister stage were up-regulated in the inoculated samples compared with the non-inoculated ones with fold changes in the range of 3.3 – 7.4 (Table 2 and S1 Fig), as were 15 genes related to translation including ribosomal proteins and elongation factors. In addition, a 16kD gamma zein was extremely up-regulated in the same comparison, making it the most highly up-regulated gene. Coincident with the up-regulation of the alpha zein genes, twelve genes coding for ribosomal proteins along with four coding for elongation factors were up-regulated at the blister stage as well. Two previous studies on transcriptional patterns in maize identified the *opaque2* transcription factor, which through a regulatory network, affects the expression of certain alpha zein proteins, together with certain ribosomal genes and elongation factors (Hunter, 2002; Li *et al.*, 2015), although DE of the *opaque2* was not detected at this stage in the current study.

In the dent stage, however, all the alpha zeins and the same gamma zein were down-regulated, the former by about 60%, and the latter by 75% (Table 4). The same pattern of down-regulation for *all* zeins was observed in the study of DGE on field inoculated ears at different stages of maturity combined (Dolezal *et al.*, 2014). In addition, two transcription factors that regulate zein expression were also down-regulated including *opaque2* (GRMZM2G015534) by 50%, and endosperm-specific prolamin box binding factor (PBF) zinc finger (GRMZM2G146283) by 60%.

An important consideration in the novel up-regulation of zeins is that TX772 has a vitreous (i.e. hard, flinty) endosperm, while most if not all of the germplasm tested in comparable DE studies are dent type (softer, more floury) endosperm (not to be confused with the dent kernel development stage). Dent types include the resistant inbred line Mp313E, which is a yellow dent type developed from Tuxpan (Scott and Zummo, 1990), and another inbred line derived from it, Mp715 (Williams and Windham, 2001). When vitreous and floury endosperms were compared for protein and starch composition, the increase in alpha zeins (twice as much in flint compared to floury) as well as the arrangement and size of starch granules contributed to the hardness of the kernel (Gayral *et al.*, 2016). Vitreous compared to softer dent type endosperm has been positively correlated with resistance to ear rot and aflatoxin contamination (Darrah *et al.*, 1987; Betrán, Isakeit and Odvody, 2002; Llorente *et al.*, 2010). Perhaps up-regulating zein genes as found at the blister stage in response to infection is one way that Tx772 builds up resistance to colonization by the fungus, as evidenced by greater expression of these genes in inoculated samples. Yet, some infection did occur in the samples harvested at the dent kernel development stage, and at that point zein gene expression in the non-inoculated kernels was two to three-fold higher than those that were inoculated. In the (Dolezal *et al.*, 2014) study, infected kernels of susceptible B73 (a softer dent kernel type) had lost much of the zein-filled hard endosperm, and with it most of the cells still capable of producing the protein, which were replaced by starchy endosperm, by maturity. In our study, most of the kernels up to the dent stage appeared to be intact, while zein gene expression was suppressed in inoculated kernels at this more advanced stage.

The second class of genes pertained to proteins that are known to increase free hexose levels, as observed by (Dolezal *et al.*, 2014). In the current study we noted up-regulation of invertase cell wall1 (GRMZM2G139300) and invertase1 (GRMZM2G394450) in the blister and dent samples, and two alpha amylase genes, (GRMZM2G103055) and (GRMZM2G138468), the latter of which was up-regulated by over 32-fold in the silk channel and side needle inoculated samples compared to the non-inoculated ones. A recent study on expression profiling of 267 unigenes in a mapping population derived from a cross between an aflatoxin contamination susceptible parent and a resistant parent revealed many genes involved in the synthesis and hydrolysis of starch and sugar mobilization were highly expressed (Dhakal *et al.*, 2017) and others related this to providing energy and/or precursors of lignin and phytoalexins used in the defense response (Bolton *et al.*, 2008; Dolezal *et al.*, 2013; Granot, David-Schwartz and Kelly, 2013; Shu *et al.*, 2015). Agrios, (2005) explained that when plants are infected by pathogens, the rate of respiration is up-regulated, which often translates to an increase in glycolysis. In more resistant plants, respiration increases more quickly to provide the abundant source of energy needed by its defense mechanisms. In the blister group, glyceraldehyde-3-phosphate dehydrogenase and an oxidoreductase, which are catalysts in the conversion of glucose to energy through their acting on NADH or NADPH, (Sirover, 2014)(Gani *et al.*, 2016) were also up-regulated by about four-fold (Table 2). In the dent development stage group, two enzymes related to cellular respiration, NADH dehydrogenase subunit 7 and protein kinase 5’AMP-activated were up-regulated as well, Table 4. Thus, our results, involving up-regulation of genes producing or utilizing simple sugars at the expense of those making starch, align with previous observations related to energy production, but not those related to the biosynthesis of lignins.

At the same time, some have suggested (Govrin and Levine, 2000; Dolezal *et al.*, 2014) that fungal pathogens alter the host plant’s metabolism to secure their own nutrition, which could certainly be the case in increasing the levels of free hexoses. In this study, fungal genes for three enzymes contributing to glycolysis were expressed in either blister or dent samples including enolase, fructose-biphosphate aldolase and glyceraldehyde 3-phosphate dehydrogenase. In addition, fungal endoglucanase was expressed, which acts as a cellulase to degrade cell walls.

### Fungal sequences show high correlation with maize DEGs and aflatoxin level

Numbers of fungal reads detected above a certain threshold (1e −20) are also listed in the Table 1 (e) and (f). Fungal cDNA sequences of mainly *A. flavus*, in addition to others identified by BLASTn (Johnson *et al.*, 2008) such as *Aspergillus orzyae*, and *Fusarium verticillioides*, were detected especially at the blister and dent stages. At the dent stage, the non-inoculated samples had what appeared to be some natural contamination from the field. Although numbers were still significantly lower than those of the inoculated samples, this is consistent with our expectation of corn development in Texas given the large pool of *A. flavus* inoculum in the field; there is currently no way to our knowledge to eliminate all natural infection under field conditions. The correlation between number of DEGs and fungal reads for each of the eight treatment groups was 0.57, but increases to 0.88 if excluding the paucity of fungal reads detected in the blister silk channel sample which had a significant number of maize DEGs. The range of aflatoxin was 0 to 137 ng g^−1^, and correlated with the distribution of fungal reads at r = .65. Levels of aflatoxin in this study (Table 1 (g)) were relatively low at the time points measured, but levels are highly subject to environmental factors (Payne and Widstrom, 2008), metabolic state of the kernels (Jiang *et al.*, 2011), the state of the fungus (Jayashree and Subramanyam, 2000), and the length of time since infection (Scott, G.E. and Zummo, 1994; Betrán and Isakeit, 2004). Mideros et al. (Mideros *et al.*, 2009) previously showed that qPCR of the *A. flavus* internal transcribed spacer 1 (ITS1) often closely correlated to aflatoxin. Our finding suggests that overall transcript level might also be a promising measure for *A. flavus* contamination.

### Characterization of fungal genes expressed

Fungal genes were identified from the samples for which at least ten reads could be assigned to a given transcript at a certain stage of maturity (S2 Table). At the milk stage, many of the fungal loci had less than ten reads, and so were not represented in this table. There were no transcripts expressed in the kernels inoculated by silk channel that were not also expressed in the side needle samples. A few fungal transcripts, especially at the dent stage were also lowly expressed in the non-inoculated samples as well as those mentioned previously. Loci of proteins or noncoding RNAs for which a specific product has not been characterized and named, were assigned to the “uncharacterized” transcripts category; these represented about 16% of total fungal transcripts.

The genes for the 40S and 60S ribosomal proteins greatly outnumbered all other genes at the blister, dough and dent stages at 58%, 8% and 36% of total transcripts at each stage respectively. At all stages, at least one stress response gene in the fungus was expressed. None of the 25 genes directly involved in the biosynthesis of the secondary metabolite aflatoxin were detected in this study (Yu *et al.*, 2004; Ehrlich, Yu and Cotty, 2005). Among the stress-related transcripts, the presence of fungal superoxide dismutase in the dent samples indicated a need reduce the levels of ROS in the kernels, which are believed to be contributory to the production of aflatoxin (Jayashree and Subramanyam, 2000; Fountain *et al.*, 2016). In addition, presence of the CpcA expressed in the dent samples was observed. CpA has been labeled a “cross pathway control” transcription factor, due to evidence that it controls transcription factors directly regulating production of fungal secondary metabolites, such as glioxin in *Aspergillus fumigatus*, or sirodesmin PL in the plant pathogen *Leptosphaeria maculans* (Desm.) (Elliott *et al.*, 2011). One other interesting gene expressed, in this case in the blister kernels was ceratO-platanin, which is an extracellular secretory protein produced during kernel colonization (Elliott *et al.*, 2011; Dolezal *et al.*, 2013). This phytotoxin elicits a response to infection that helps establish and maintain disease in the host plant.

## Conclusion

RNA-Seq with *de novo* transcriptome assembly in Trinity served to illuminate different patterns of differential expression among the four stages of maturity in maize kernels and identify the DE of many genes in maize kernels in response to field inoculation with *Aspergillus flavus*. Both the silk channel (non-wounding) and the side-needle technique (wounding) were effective in establishing *A. flavus* fungal infections as evidenced by the detection of fungal reads in inoculated samples and similar DGE in magnitude and direction at each stage of maturity, even with the limited number of replications. Sixteen of the DGEs identified had been previously associated with a resistance response to the presence of a pathogen or tissue damage, and certain others pertained to carbohydrate metabolism and energy production to support the defense response. Any future work in differential gene expression should critically consider the development stage of the seed when evaluating significant differences among genotypes or treatments. In the case of flint endosperm types in maize, we have provided some evidence of the contribution of alpha zeins can make to resistance to infection and levels of mycotoxin contamination. The complexity of biology and especially gene network analysis means that a single study can often not be definitive, and a body of evidence must be built, so it is important that here we both confirmed 16 previously implicated genes and identified additional genetic pathways for future investigation.

Testing this further, a more extensive DGE study beyond this exploratory one is recommended, preferably on endosperm tissue alone collected at two or three time points with more replications. This would provide opportunity to confirm that up-regulation of zein genes in the early stages is a novel feature of germplasm such as Tx772. Another modification would be to select only one method of inoculation and apply that without the fungal suspension for mock-inoculated controls. And finally, valuable information would be gained from expanding the same tests to include a well-known susceptible inbred such as B73 or Va35. The results would indicate which genes are most likely to contribute to aflatoxin resistance in the same environments, and enable more direct comparisons with the results of similar studies.

## Acknowledgments

The authors would like to thank the following contributors to this study: Jingjia Li, a graduate student in Hong-Bin Zhang’s laboratory who greatly assisted with the library preparation for sequencing, and Crystal Buchanan at HPRC for technical advice. We are also most grateful for the expert advice regarding the application of Trinity for *de novo* assembly of our RNA-Seq data to Brian Haas, the primary developer of Trinity at The Broad Institute at MIT, Mark Chapman of the University of Southhampton, UK, Ken Field of Bicknell University and Tiago Hori of The Center for AquaCulture Technologies.

Graduate study was supported by USDA National Institute of Food and Agriculture Competitive Grant 2014-6804-21836.

## Supplemental Data Files

**S1 Table. Differentially expressed genes associated with resistance to pathogens**

**S2 Table. Annotation of fungal genes expressed in maize kernels at different stages of maturity**

**S1 Figure. Normalized read counts of α zein genes at different stages of maturity and under different treatments**

## References

Afzal, A. J., Wood, A. J. and Lightfoot, D. A. (2008) ‘Plant Receptor-Like Serine Threonine Kinases: Roles in Signaling and Plant Defense’, Molecular Plant-Microbe Interactions, 21(5), pp. 507–517. doi: 10.1094/mpmi-21-5-0507.

Agrios, G. N. (2005) Plant Pathology. 5th edn. Burlington, MA: Elsevier Ltd.

Apweiler, R. et al. (2014) ‘Activities at the Universal Protein Resource (UniProt)’, Nucleic Acids Research, 42(D1), pp. 191–198. doi: 10.1093/nar/gkt1140.

Benjamini, Y. and Hochberg, Y. (2018) ‘Controlling the False Discovery Rate: A Practical and Powerful Approach to Multiple Testing’, Journal of the Royal Statistical Society: Series B (Methodological), 57(1), pp. 289–300. doi: 10.1111/j.2517-6161.1995.tb02031.x.

Betrán, F. J. and Isakeit, T. (2004) ‘Aflatoxin accumulation in maize hybrids of different maturities’, Agronomy Journal, 96(2), pp. 565–570.

Betrán, F. J., Isakeit, T. and Odvody, G. (2002) ‘Aflatoxin accumulation of white and yellow maize inbreds in diallel crosses’, Crop Science, 42(6), pp. 1894–1901.

Bolton, M. D. et al. (2008) ‘Lr34-Mediated Leaf Rust Resistance in Wheat: Transcript Profiling Reveals a High Energetic Demand Supported by Transient Recruitment of Multiple Metabolic Pathways’, Molecular Plant-Microbe Interactions, 21(12), pp. 1515–1527. doi: 10.1094/mpmi-21-12-1515.

Brueggeman, R. et al. (2002) ‘The barley stem rust-resistance gene Rpg1 is a novel disease-resistance gene with homology to receptor kinases’, PNAS, 99(14), pp. 9328–33.

Buckley, P., Williams, W. and Windham, G. (2006) Publication: USDA ARS, USDA ARS PUB 179111. Available at: https://www.ars.usda.gov/research/publications/publication/?seqNo115=179111 (Accessed: 27 March 2019).

Chen, T. et al. (2016) ‘Two members of TaRLK family confer powdery mildew resistance in common wheat’, BMC Plant Biology. BMC Plant Biology, 16(1), pp. 1–17. doi: 10.1186/s12870-016-0713-8.

Christensen, S. A. et al. (2013) ‘The maize lipoxygenase, ZmLOX10, mediates green leaf volatile, jasmonate and herbivore-induced plant volatile production for defense against insect attack’, Plant Journal, 74(1), pp. 59–73. doi: 10.1111/tpj.12101.

Cordes, M. et al. (2009) ‘The B73 Maize Genome: Complexity, Diversity, and Dynamics’, Science, 326(5956), pp. 1112–1115. doi: 10.1126/science.1178534.

Darrah, L. L. et al. (1987) ‘Inheritance of aflatoxin B1 levels in maize kernels under modified natural inoculation with Aspergillus flavus’, Crop Science, 27, pp. 869–872.

DeYoung, B. J. and Innes, R. W. (2006) ‘Plant NBS-LRR proteins in pathogen sensing and host defense’, Nature Immunology, 7(12), pp. 1243–1249. doi: 10.1038/ni1410.

Dhakal, R. et al. (2017) ‘Expression Profiling Coupled with In-silico Mapping Identifies Candidate Genes for Reducing Aflatoxin Accumulation in Maize’, Frontiers in Plant Science. Frontiers, 8, pp. 1–15. doi: 10.3389/fpls.2017.00503.

Dillies, M.-A. et al. (2013) ‘A comprehensive evaluation of normalization methods for Illumina high-throughput RNA sequencing data analysis’, Briefings in Bioinformatics. Narnia, 14(6), pp. 671–683. doi: 10.1093/bib/bbs046.

Dolezal, A. L. et al. (2013) ‘Localization, morphology and transcriptional profile of Aspergillus flavus during seed colonization’, Molecular Plant Pathology, 14(9), pp. 898–909. doi: 10.1111/mpp.12056.

Dolezal, A. L. et al. (2014) ‘Aspergillus flavus infection induces transcriptional and physical changes in developing maize kernels’, Frontiers in Microbiology, 5(July), pp. 1–10. doi: 10.3389/fmicb.2014.00384.

Ebrahim, S., Usha, K. and Singh, B. (2011) ‘Pathogenesis related (PR) proteins in plant defense mechanism’, Sci Against Microb Pathog, 2, pp. 1043–1054.

Ehrlich, K. C., Yu, J. and Cotty, P. J. (2005) ‘Aflatoxin biosynthesis gene clusters and flanking regions’, Journal of Applied Microbiology, 99(3), pp. 518–527. doi: 10.1111/j.1365-2672.2005.02637.x.

Elliott, C. E. et al. (2011) ‘The cross-pathway control system regulates production of the secondary metabolite toxin, sirodesmin PL, in the ascomycete, Leptosphaeria maculans’, BMC Microbiology. BioMed Central Ltd, 11(1), p. 169. doi: 10.1186/1471-2180-11-169.

Fajardo, J. E. et al. (1995) ‘Phenolic compounds in peanut seeds: Enhanced elicitation by chitosan and effects on growth and aflatoxin B 1 production by *Aspergillus flavus*’, Food Biotechnology. Taylor & Francis Group, 9(1–2), pp. 59–78. doi: 10.1080/08905439509549885.

Fajardo, J. E. et al. (2009) ‘Effects of chitosan and aspergillus flaws on isozymes related to phenouc compound synthesis and protein profiles of peanut seeds’, Food Biotechnology, 8(2–3), pp. 213–228. doi: 10.1080/08905439409549876.

Farfan, I. D. B. et al. (2015) ‘Genome wide association study for drought, aflatoxin resistance, and important agronomic traits of maize hybrids in the sub-tropics’, PLoS ONE, 10(2), pp. 1–30. doi: 10.1371/journal.pone.0117737.

Ferreira, R. B. et al. (2007) ‘The role of plant defence proteins in fungal pathogenesis’, Molecular Plant Pathology. John Wiley & Sons, Ltd (10.1111), 8(5), pp. 677–700. doi: 10.1111/j.1364-3703.2007.00419.x.

Finn, R. D. et al. (2014) ‘Pfam: the protein families database’, Nucleic Acids Research. Narnia, 42(D1), pp. D222–D230. doi: 10.1093/nar/gkt1223.

Fountain, J. C. et al. (2014) ‘Environmental influences on maize-Aspergillus flavus interactions and aflatoxin production’, Frontiers in Microbiology. Frontiers, 5, p. 40. doi: 10.3389/fmicb.2014.00040.

Fountain, J. C. et al. (2016) ‘Responses of Aspergillus flavus to oxidative stress are related to fungal development regulator, antioxidant enzyme, and secondary metabolite biosynthetic gene expression’, Frontiers in Microbiology, 7(DEC), pp. 1–16. doi: 10.3389/fmicb.2016.02048.

Gani, Z. et al. (2016) ‘Purification and characterization of glyceraldehyde-3-phosphate-dehydrogenase (GAPDH) from pea seeds’, Protein Expression and Purification. Academic Press, 127, pp. 22–27. doi: 10.1016/J.PEP.2016.06.014.

Gayral, M. et al. (2016) ‘Transition from vitreous to floury endosperm in maize (Zea mays L.) kernels is related to protein and starch gradients’, Journal of Cereal Science, 68, pp. 148–154. doi: 10.1016/j.jcs.2016.01.013.

Glazebrook, J. (2005) ‘Contrasting Mechanisms of Defense Against Biotrophic and Necrotrophic Pathogens’, Annual Review of Phytopathology, 43(1), pp. 205–227. doi: 10.1146/annurev.phyto.43.040204.135923.

Goodman, M. M. (2005) ‘Broadening the U.S. maize germplasm base’, Maydica, 50(3–4), pp. 203–214.

Govrin, E. M. and Levine, A. (2000) ‘The hypersensitive response facilitates plant infection by the necrotrophic pathogen Botrytis cinerea’, Current Biology, 10(13), pp. 751–757.

Granot, D., David-Schwartz, R. and Kelly, G. (2013) ‘Hexose Kinases and Their Role in Sugar-Sensing and Plant Development’, Frontiers in Plant Science, 4(March), pp. 1–17. doi: 10.3389/fpls.2013.00044.

Grubisha, L. C. and Cotty, P. J. (2015) ‘Genetic Analysis of the Aspergillus flavus Vegetative Compatibility Group to Which a Biological Control Agent That Limits Aflatoxin Contamination in U.S. Crops Belongs.’, Applied and environmental microbiology. American Society for Microbiology, 81(17), pp. 5889–99. doi: 10.1128/AEM.00738-15.

Haas, B. J. et al. (2013) ‘De novo transcript sequence reconstruction from RNA-seq using the Trinity platform for reference generation and analysis’, Nature Protocols. Nature Publishing Group, 8(8), pp. 1494–1512. doi: 10.1038/nprot.2013.084.

Hawkins, L. K. et al. (2015) ‘Characterization of the maize chitinase genes and their effect on Aspergillus flavus and aflatoxin accumulation resistance’, PLoS ONE, 10(6). doi: 10.1371/journal.pone.0126185.

Hunter, B. G. (2002) ‘Maize Opaque Endosperm Mutations Create Extensive Changes in Patterns of Gene Expression’, the Plant Cell Online, 14(10), pp. 2591–2612. doi: 10.1105/tpc.003905.

Isakeit, T. et al. (2010) ‘Biological control of aflatoxin contamination of corn in Texas using Afla-Guard (R), a commercial atoxigenic strain of Aspergillus flavus’, Canadian Journal of Plant Pathology-Revue Canadienne De Phytopathologie, 32(3), pp. 407–408.

Jayashree, T. and Subramanyam, C. (2000) ‘Oxidative stress as a prerequisite for aflatoxin production by Aspergillus parasiticus’, Free Radical Biology and Medicine, 29(10), pp. 981–985. doi: 10.1016/S0891-5849(00)00398-1.

Jiang, T. et al. (2011) ‘Expression analysis of stress-related genes in kernels of different maize (zea mays l.) inbred lines with different resistance to aflatoxin contamination’, Toxins, 3(6), pp. 538–550. doi: 10.3390/toxins3060538.

Johnson, M. et al. (2008) ‘NCBI BLAST: a better web interface’, Nucleic Acids Research. Narnia, 36(Web Server), pp. W5–W9. doi: 10.1093/nar/gkn201.

Kelley, R. Y. et al. (2012) ‘Identification of Maize Genes Associated with Host Plant Resistance or Susceptibility to Aspergillus flavus Infection and Aflatoxin Accumulation’, PLoS ONE. Edited by G. G. Vendramin. Public Library of Science, 7(5), p. e36892. doi: 10.1371/journal.pone.0036892.

Lawrence, C. J. et al. (2008) ‘MaizeGDB: The maize model organism database for basic, translational, and applied research.’, International journal of plant genomics. Hindawi, 2008, pp. 1–10. doi: 10.1155/2008/496957.

Lay, F. T. et al. (2003) ‘The three-dimensional solution structure of NaD1, a new floral defensin from Nicotiana alata and its application to a homology model of the crop defense protein alfAFP’, Journal of Molecular Biology, 325(1), pp. 175–188. doi: 10.1016/S0022-2836(02)01103-8.

Lee, J. M. et al. (2002) ‘DNA array profiling of gene expression changes during maize embryo development’, Functional and Integrative Genomics, 2(1–2), pp. 13–27. doi: 10.1007/s10142-002-0046-6.

Li, C. et al. (2015) ‘Genome-Wide Characterization of cis-Acting DNA Targets Reveals the Transcriptional Regulatory Framework of Opaque2 in Maize OPEN’, Plant Cell, 27(3), pp. 532–45. doi: 10.1105/tpc.114.134858.

Liang, X., Luo, M. and Guo, B. (2006) ‘Resistance mechanisms to Aspergillus flavus infection and aflatoxin contamination in peanut (Arachis hypogaea)’, Plant Pathology Journal, 5(1), pp. 115–24.

Libault, M. et al. (2007) ‘Identification of 118 Arabidopsis Transcription Factor and 30 Ubiquitin-Ligase Genes Responding to Chitin, a Plant-Defense Elicitor’, Molecular Plant-Microbe Interactions, 20(8), pp. 900–911. doi: 10.1094/mpmi-20-8-0900.

Liu, J. et al. (2012) ‘Genome-wide analysis uncovers regulation of long intergenic noncoding RNAs in Arabidopsis.’, The Plant cell. American Society of Plant Biologists, 24(11), pp. 4333–45. doi: 10.1105/tpc.112.102855.

Liu, X. et al. (2008) ‘Genome-wide analysis of gene expression profiles during the kernel development of maize (Zea mays L.)’, Genomics, 91(4), pp. 378–387. doi: 10.1016/j.ygeno.2007.12.002.

Liu, Y. et al. (2004) ‘Molecular chaperone Hsp90 associates with resistance protein N and its signaling proteins SGT1 and Rar1 to modulate an innate immune response in plants.’, The Journal of biological chemistry. American Society for Biochemistry and Molecular Biology, 279(3), pp. 2101–8. doi: 10.1074/jbc.M310029200.

Llorente, C. F. et al. (2010) ‘Registration of Tx772 Maize’, Crop Science, 44(3), p. 1036–a. doi: 10.2135/cropsci2004.1036a.

Lu, X. et al. (2013) ‘The Differential Transcription Network between Embryo and Endosperm in the Early Developing Maize Seed 1[C][W][OA]’, Plant Physiology Ò, 162, pp. 440–455. doi: 10.1104/pp.113.214874.

Marsh, S. F. and Payne, G. A. (1984) ‘Preharvest infection of corn silks and kernels by Aspergillus flavus’, Ecology and Epidemiology, 74, pp. 1284–1289. Available at: https://www.apsnet.org/publications/phytopathology/backissues/Documents/1984Articles/Phyto74n11_1284.pdf (Accessed: 27 March 2019).

Mideros, S. X. et al. (2009) ‘Aspergillus flavus Biomass in Maize Estimated by Quantitative Real-Time Polymerase Chain Reaction Is Strongly Correlated with Aflatoxin Concentration’, Plant Disease, 93(11), pp. 1163–1170. doi: 10.1094/pdis-93-11-1163.

Mideros, S. X. et al. (2014) ‘Quantitative Trait Loci Influencing Mycotoxin Contamination of Maize: Analysis by Linkage Mapping, Characterization of Near-Isogenic Lines, and Meta-Analysis’, Crop Science. The Crop Science Society of America, Inc., 54(1), p. 127. doi: 10.2135/cropsci2013.04.0249.

Mittler, R. et al. (1999) ‘Transgenic tobacco plants with reduced capability to detoxify reactive oxygen intermediates are hyperresponsive to pathogen infection.’, Proceedings of the National Academy of Sciences of the United States of America. National Academy of Sciences, 96(24), pp. 14165–70. doi: 10.1073/PNAS.96.24.14165.

Monaco, M. K. et al. (2013) ‘Maize Metabolic Network Construction and Transcriptome Analysis’, The Plant Genome. Crop Science Society of America, 6(1), p. 0. doi: 10.3835/plantgenome2012.09.0025.

Naumann, T. A. and Wicklow, D. T. (2010) ‘Allozyme-Specific Modification of a Maize Seed Chitinase by a Protein Secreted by the Fungal Pathogen Stenocarpella maydis’, Phytopathology, 100(7), pp. 645–654. doi: 10.1094/phyto-100-7-0645.

Naumann, T. A., Wicklow, D. T. and Kendra, D. F. (2009) ‘Maize seed chitinase is modified by a protein secreted by Bipolaris zeicola’, Physiological and Molecular Plant Pathology. Elsevier Ltd, 74(2), pp. 134–141. doi: 10.1016/j.pmpp.2009.10.004.

Odvody, G. et al. (2000) Aflatoxin and insect respones of near-isogenic Bt and non-Bt commercial corn hybrids in south Texas. Fish Camp, CA.

Ogunola, O. F. et al. (2017) ‘Characterization of the maize lipoxygenase gene family in relation to aflatoxin accumulation resistance’, PLOS ONE. Edited by P. L. Kulwal. Public Library of Science, 12(7), p. e0181265. doi: 10.1371/journal.pone.0181265.

Payne, G. A. and Widstrom, N. W. (2008) ‘Aflatoxin in maize’, Critical Reviews in Plant Sciences, 10(5), pp. 423–440. doi: 10.1080/07352689209382320.

Reese, B. N. et al. (2011) ‘Gene Expression Profile and Response to Maize Kernels by Aspergillus flavus’, Phytopathology, 101(7), pp. 797–804. doi: 10.1094/phyto-09-10-0261.

Ritchie, S. W., Hanway, J. J. and Benson, G. O. (1997) How a corn plant develops.; Spec. Publ. 48. Ames, IA.

Sawano, Y. et al. (2007) ‘Purification, characterization, and molecular gene cloning of an antifungal protein from Ginkgo biloba seeds’, Biological Chemistry, 388(3), pp. 273–280. doi: 10.1515/BC.2007.030.

Scott, G.E. and Zummo, N. (1994) ‘Kernel infection and aflatoxin prod in maize by A flavus rel to inoc and harvest.pdf’, Plant Disease, 78, pp. 123–125.

Scott, G. E. et al. (1991) ‘Aflatoxin in Corn Hybrids Field Inoculated with Aspergillus flavus’, Agronomy Journal. American Society of Agronomy, 83(3), p. 595. doi: 10.2134/agronj1991.00021962008300030018x.

Scott, G. E. and Zummo, N. (1990) ‘Registration of Mp313E parental line of maize’, Crop Science, 30(6), p. 1378.

Shadle, G. et al. (2007) ‘Down-regulation of hydroxycinnamoyl CoA: Shikimate hydroxycinnamoyl transferase in transgenic alfalfa affects lignification, development and forage quality’, Phytochemistry, 68(11), pp. 1521–1529. doi: 10.1016/j.phytochem.2007.03.022.

Shigeoka, S. et al. (2002) ‘Regulation and function of ascorbate peroxidase isoenzymes’, Journal of Experimental Botany. Narnia, 53(372), pp. 1305–1319. doi: 10.1093/jexbot/53.372.1305.

Shu, X. et al. (2015) ‘Tissue-specific gene expression in maize seeds during colonization by *Aspergillus flavus* and *Fusarium verticillioides*’, Molecular Plant Pathology. John Wiley & Sons, Ltd (10.1111), 16(7), pp. 662–674. doi: 10.1111/mpp.12224.

Sirover, M. A. (2014) ‘Structural analysis of glyceraldehyde-3-phosphate dehydrogenase functional diversity’, International Journal of Biochemistry and Cell Biology. Elsevier Ltd, 57, pp. 20–26. doi: 10.1016/j.biocel.2014.09.026.

Smith, S. D., Murray, S. C. and Heffner, E. (2015) ‘Molecular analysis of genetic diversity in a Texas maize (Zea mays L) breeding program’, Maydica, 60(2).

Solomon, M. et al. (1999) ‘The Involvement of Cysteine Proteases and Protease Inhibitor Genes in the Regulation of Programmed Cell Death in Plants’, the Plant Cell Online, 11(3), pp. 431–444. doi: 10.1105/tpc.11.3.431.

Tang, Y. et al. (2015) ‘Genome-Wide Characterization of cis-Acting DNA Targets Reveals the Transcriptional Regulatory Framework of Opaque2 in Maize’, The Plant Cell, 27(3), pp. 532–545. doi: 10.1105/tpc.114.134858.

Vignols, F. et al. (2007) ‘The brown midrib3 (bm3) Mutation in Maize Occurs in the Gene Encoding Caffeic Acid O-Methyltransferase’, The Plant Cell, 7(4), p. 407. doi: 10.2307/3870079.

Wahl, N. et al. (2017) ‘Identification of Resistance to Aflatoxin Accumulation and Yield Potential in Maize Hybrids in the Southeast Regional Aflatoxin Trials (SERAT)’, Crop Science. The Crop Science Society of America, Inc., 57(1), p. 202. doi: 10.2135/cropsci2016.06.0519.

Warburton, M. L. et al. (2013) ‘Phenotypic and Genetic Characterization of a Maize Association Mapping Panel Developed for the Identification of New Sources of Resistance to and Aflatoxin Accumulation’, Crop Science. The Crop Science Society of America, Inc., 53(6), p. 2374. doi: 10.2135/cropsci2012.10.0616.

Williams, W. P. and Windham, G. L. (2001) ‘Registration of maize germplasm line Mp715’, Crop Science, 41(4), pp. 1374–1375.

Wisser, R. J. et al. (2011) ‘Multivariate analysis of maize disease resistances suggests a pleiotropic genetic basis and implicates a GST gene’, PNAS, 108(18), pp. 7339–44. doi: 10.1073/pnas.1011739108.

Woo, Y.-M. et al. (2001) Genomics Analysis of Genes Expressed in Maize Endosperm Identifies Novel Seed Proteins and Clarifies Patterns of Zein Gene Expression, The Plant Cell. Available at: www.plantcell.org (Accessed: 27 March 2019).

Xu, Z.-S. et al. (2012) ‘Heat Shock Protein 90 in Plants: Molecular Mechanisms and Roles in Stress Responses’, International Journal of Molecular Sciences. Multidisciplinary Digital Publishing Institute, 13(12), pp. 15706–15723. doi: 10.3390/ijms131215706.

Yu, J. et al. (2004) ‘Clustered Pathway Genes in Aflatoxin Biosynthesis’, Appl. Environ. Microbiol. American Society for Microbiology, 70(3), pp. 1253–1262. doi: 10.1128/AEM.70.3.1253-1262.2004.

Zhang, B. et al. (2007) ‘Characterization of Early, Chitin-Induced Gene Expression in Arabidopsis’, Molecular Plant-Microbe Interactions, 15(9), pp. 963–970. doi: 10.1094/mpmi.2002.15.9.963.

Zhang, L. et al. (2009) ‘Identification of an Apoplastic Protein Involved in the Initial Phase of Salt Stress Response in Rice Root by Two-Dimensional Electrophoresis 1[C][W][OA]’, Plant Physiology, 149(2), pp. 916–28. doi: 10.1104/pp.108.131144.

Zummo, N. and Scott, G. E. (1989) ‘Evaluation of Field Inoculation Techniques for Screening Maize Genotypes Against Kernel Infection by Aspergillus-Flavus in Mississippi Usa’, Plant Disease, pp. 313–316.

