## Supplementary material for "Maize Kernel Development Stage the Primary Factor in Differential Gene Expression in Response to Two Methods of Field Inoculation with *Aspergillus flavus*": S1 Figure (from another version of this preprint: version 1, 2019-04-26)

|  | α zein 1 | α zein 2 | α zein 3 | α zein 4 | α zein 5 | α zein 6 | α zein 7 | α zein 8 |
| --- | --- | --- | --- | --- | --- | --- | --- | --- |
| AD | 1198 | 21104 | 10111 | 2976 | 1531 | 33338 | 5316 | 10692 |
| AK | 1314 | 25043 | 11605 | 2356 | 1791 | 41021 | 6134 | 8791 |
| AN | 189 | 3717 | 1636 | 673 | 342 | 9709 | 850 | 2402 |
| BD1 | 761 | 13684 | 7888 | 2822 | 1273 | 24952 | 4325 | 5942 |
| BD2 | 739 | 15420 | 7383 | 2053 | 1091 | 26317 | 3959 | 7192 |
| BD3 | 649 | 11232 | 5853 | 2555 | 1112 | 18979 | 3938 | 6019 |
| BK1 | 495 | 9962 | 5414 | 2002 | 1140 | 14138 | 3496 | 5495 |
| BK2 | 597 | 12087 | 5350 | 2417 | 1195 | 18388 | 3848 | 6756 |
| BN1 | 600 | 13160 | 7163 | 2275 | 1153 | 20572 | 3435 | 6120 |
| BN2 | 670 | 13759 | 8126 | 2648 | 1440 | 21846 | 4854 | 7823 |
| ED1 | 152 | 3607 | 1377 | 1004 | 186 | 5535 | 837 | 1843 |
| ED2 | 279 | 5120 | 2419 | 1835 | 569 | 8054 | 1811 | 2488 |
| EK1 | 163 | 3776 | 1524 | 1147 | 340 | 4544 | 1174 | 1746 |
| EK2 | 124 | 3025 | 1479 | 1015 | 314 | 4374 | 910 | 1718 |
| EN1 | 177 | 4727 | 1966 | 1089 | 381 | 6013 | 1690 | 2383 |
| EN2 | 257 | 5417 | 2254 | 1635 | 385 | 8419 | 1422 | 2156 |
| FD1 | 61 | 1169 | 479 | 508 | 85 | 1610 | 332 | 740 |
| FD2 | 78 | 1203 | 669 | 871 | 91 | 2128 | 418 | 986 |
| FK1 | 72 | 1548 | 652 | 712 | 116 | 2396 | 454 | 881 |
| FK2 | 76 | 1631 | 889 | 864 | 214 | 2605 | 604 | 1540 |
| FN1 | 80 | 1806 | 821 | 786 | 155 | 2763 | 684 | 1414 |
| FN2 | 179 | 3917 | 1657 | 1051 | 373 | 5652 | 1332 | 2343 |

S1 Figure. Normalized read counts of α zein genes at different stages of maturity and under different treatments
