## Supplementary material for "Maize Kernel Development Stage the Primary Factor in Differential Gene Expression in Response to Two Methods of Field Inoculation with *Aspergillus flavus*": S1 Table (from another version of this preprint: version 1, 2019-04-26)

| **S1 Table. Differentially expressed genes associated with resistance to pathogens** | | | | | |
| --- | --- | --- | --- | --- | --- |
| **References** | | **Gene** | | **Gramene #** | **Biological Process** |
| 4, 5, 7 | | Chitinase family 19 | | GRMZM2G005633 | antifungal, degrades fungal cell walls |
| 4, 5, 7 | | Chitinase family 18 | | GRMZM2G057093 | antifungal, degrades fungal cell walls |
| 3, 18 | | Lipoxygenase 1 (LOX1) | | GRMZM2G156861 | stress or wound induced |
| 3, 18 | | Lipoxygenase 8 (LOX8) | | GRMZM5G822593 | stress or wound induced |
| 5 | | Flower-specific γ-thionine | | GRMZM2G392863 | fungal inhibition through ion efflux mechanism |
| 9, 13 | | Protein kinase (related to salt stress) | | GRMZM2G334181 | antifungal |
| 11, 16, 17 | | Cysteine protease | | GRMZM2G150256 | hypersensitive response |
| 4, 6, 14 | | Ascorbate peroxidase | | GRMZM2G140970 | strong antioxidant, removes hydrogen peroxide |
| 2, 18 | | Jasmonate-induced protein | | GRMZM2G050412 | stress response, disease resistance |
| 2, 18 | | Beta glucosidase jasmonate-induced aggregating factor 1 | | GRMZM2G172204 | stress response, disease resistance |
| 8, 13, 19 | | Glutathione-S-transferases (GST) | | GRMZM2G042639 | detoxification of toxic substances |
| 10, 20 | | Hsp90 | | GRMZM2G112165 | reduce plant stress, molecular chaperone |
| 1, 9 | | Receptor-like ser/thr kinases with LRR | | GRMZM2G011526 | signaling in pathogen recognition |
| 12 | | Bzip transcription factor 60 | | GRMZM2G444748 | transcription factor regulating response to chitin |
| 10, 13 | | Hydroxycinnamoyl transferase | | GRMZM2G127251 | lignin biosynthesis |
| 10, 13 | | O-methyltransferase | | GRMZM2G036048 | lignin biosynthesis |
| **References for genes and/or metabolic pathways** | | | | | |
| 1 | Afzal et al., 2008 | | Plant receptor-like serine threonine kinases: roles in signaling and plant defense | | |
| 2 | Chaudry et al., 1994 | | The barley 60 kDA jasmonate-induced protein (JIP60) is a novel ribosome-inactivating protein | | |
| 3 | Christensen et al., 2013 | | The maize liposygenase, ZmLOX10, mediates green leaf volatile, jasmonate and herbivore-induced plant volatile production for defense against insect attack | | |
| 4 | Dolezal et al., 2015 | | Aspergillus flavus infection induces transcriptional and physical changes in developing maize kernels | | |
| 5 | Ferreira et al., 2007 | | The role of plant defense proteins in fungal pathogenesis | | |
| 6 | Fortunato et al., 2015 | | Changes in the antioxidant system in soybean leaves infected by Corynespora cassiicola | | |
| 7 | Hawkins et al., 2015 | | Characterization of the maize chitinase genes and their effect on Aspergillus flavus and aflatoxin accumulation resistance | | |
| 8 | Holt et al., 1995 | | Characterization of the safener-induced glutathione S-transferase isoform II from maize | | |
| 9 | Jiang et al., 2011 | | Expression analysis of stress-related genes in kernels of different maize (Zea mays L.) inbred lines with different resistance to aflatoxin contamination | | |
| 10 | Kelley et al., 2012 | | Identification of maize genes associated with host plant resistance or susceptibility to Aspergillus flavus infection and aflatoxin | | |
| 11 | Lampl et al., 2013 | | Set-point of RD21 protease activity by ATSerpin1 controls cell death in Arabidopsis | | |
| 12 | Libault et al., 2007 | | Identification of 118 Arabidopsis transcription factor and 30 ubiquitin-ligase genes responding to chitin, a plant-defense elicitor | | |
| 13 | Luo et al., 2011 | | Transcriptional profiles uncover Aspergillus flavus-induced resistance in maize kernels | | |
| 14 | Pechanova et al., 2011 | | Proteomic analysis of the maize rachis: Potential roles of constitutive and induced proteins in resistance to | | |
| 15 | Shigeoka et al., 2002 | | Regulation and function of ascorbate peroxidase isoenzymes | | |
| 16 | Shindo et al., 2012 | | A role in immunity for Arabidopsis cysteine protease RD21, the ortholog of the tomato immune protease C14 | | |
| 17 | Solomon et al., 1999 | | The involvement of cysteine proteases and protease inhibitor genes in the regulation of programmed cell death in plants | | |
| 18 | Tang et al., 2015 | | Using genome-wide associations to identify metabolic pathways involved in maize aflatoxin accumulation resistance | | |
| 19 | Wisser et al., 2011 | | Multivariate analysis of maize disease resistances suggests a pleiotropic genetic basis and implicates a GST gene | | |
| 20 | Xu et al., 2012 | | Heat Shock Protein 90 in Plants: Molecular Mechanisms and Roles in Stress Responses | | |
