## Supplementary material for "Maize Kernel Development Stage the Primary Factor in Differential Gene Expression in Response to Two Methods of Field Inoculation with *Aspergillus flavus*": S2 Table (from another version of this preprint: version 1, 2019-04-26)

| **S2 Table. Annotation of fungal genes expressed in maize kernels at different stages of maturity** | | | | | |
| --- | --- | --- | --- | --- | --- |
| **Identified Fungal Genes** | **Blister** | **Milk** | **Dough** | **Dent** | **Annotation** |
| ATP synthase |  |  |  |  | ATP synthesis |
| fucose-specific lectin FleA |  |  |  |  | carbohydrate binding/human disease |
| 1,3-beta-glucanosyltransferase |  |  |  |  | carbohydrate metabolism |
| endoglucanse |  |  |  |  | carbohydrate metabolism |
| enolase/allergen Asp F 22 |  |  |  |  | carbohydrate metabolism |
| fructose-bisphosphate aldolase |  |  |  |  | carbohydrate metabolism |
| glyceraldehyde 3-phos dehydrogenase |  |  |  |  | carbohyrate metabolism |
| plasma membrane H+-ATPase |  |  |  |  | cell metabolism |
| thiamine biosynthesis protein |  |  |  |  | cell metabolism |
| thiozole biosynthesis enzyme |  |  |  |  | cell metabolism |
| alcohol dehydrogenase |  |  |  |  | cell metabolism / detox of alcohol |
| aldehyde dehydrogenase |  |  |  |  | cell metabolism / detox of aldehyde |
| cytochrome c oxidase |  |  |  |  | cellular respiration |
| extracellular 3-ketosteroid 1-dehydrogenase |  |  |  |  | cellular respiration |
| mitochondrial F1 ATPase subunit alpha |  |  |  |  | cellular respiration |
| pyruvate decarboxylase |  |  |  |  | cellular respiration (anaerobic) |
| FKBP-type peptidyl prolyl isomerase |  |  |  |  | chaperone activity |
| histone H3 |  |  |  |  | DNA replication, repair, transcription |
| histone H4 |  |  |  |  | DNA replication, repair, transcription |
| histone H4.1 |  |  |  |  | DNA replication, repair, transcription |
| woronin body major protein precursor |  |  |  |  | fungal hyphae protection |
| Bax Inhibitor family protein |  |  |  |  | ion transport regulation |
| allergenic cerat0-platanin |  |  |  |  | pathogenesis / elicitor of host response |
| allergen Asp F3 |  |  |  |  | pathogenesis / allergenic response |
| pectinesterase precursor |  |  |  |  | pathogenesis / cell wall degradation |
| polyubiquitin UbiD/Ubi4 |  |  |  |  | protein degradation |
| Glutamate/Leucine/Phenylalanine/Valine dehydrogenase |  |  |  |  | protein metabolism |
| glutamine synthetase |  |  |  |  | protein metabolism |
| peptidyl-prolyl cis-trans isomerase |  |  |  |  | protein metabolism/chaperone |
| Ribosomal Protein 40S |  |  |  |  | protein synthesis |
| Ribosomal Protein 60S |  |  |  |  | protein synthesis |
| transl elong factor eEF-3 |  |  |  |  | protein synthesis |
| transl elong factor EF-1 |  |  |  |  | protein synthesis |
| transl init factor eIF-5A |  |  |  |  | protein synthesis |
| translation initiation factor 4 |  |  |  |  | protein synthesis |
| nucleoside diphosphate kinase |  |  |  |  | signal transduction |
| phosphoglycerate kinase PgkA |  |  |  |  | signal transduction |
| Aha1 domain family |  |  |  |  | stress response |
| CipC-like antibiotic stress responsive protein |  |  |  |  | stress response |
| flavohemoprotein |  |  |  |  | stress response |
| Hsp12 |  |  |  |  | stress response |
| HSP30/HSP42 |  |  |  |  | stress response |
| Hsp30-like |  |  |  |  | stress response |
| Hsp70 |  |  |  |  | stress response |
| Hsp90/Hsp1 |  |  |  |  | stress response |
| Mod-E/Hsp90/Hsp1 |  |  |  |  | stress response |
| superoxide dismutase |  |  |  |  | stress response |
| actin Act1 |  |  |  |  | structural |
| GPI-anchored cell wall org protein |  |  |  |  | structural |
| anchored serine-rich protein |  |  |  |  | structural |
| bZIP transcription factor CpcA |  |  |  |  | transcription regulation |
| BYS1 domain protein |  |  |  |  | unknown function |
| extracell serine-threonine rich protein |  |  |  |  | unknown function |
| high expression lethality protein Hel10 |  |  |  |  | unknown function |
| Uncharacterized |  |  |  |  | uncharacterized |
| Number of Unique Transcripts Detected (No. reads ≥ 10) | 70 | 7 | 25 | 100 |  |
| Key to chart: Med blue: side needle only, Red: side needle and |  |  |  |  |  |
| silk channel, Light blue: side needle, silk channel and noninoculated |  |  |  |  |  |
